# Eusocial insect queens show costs of reproduction and transcriptomic signatures of reduced longevity

**DOI:** 10.1101/2022.03.20.485028

**Authors:** David H. Collins, David C. Prince, Jenny L. Donelan, Tracey Chapman, Andrew F. G. Bourke

## Abstract

Eusocial insect queens have been suggested to be counter-examples to the standard evolutionary theory of ageing through lacking costs of reproduction. Using the bumblebee (*Bombus terrestris*), we tested this hypothesis experimentally against the alternative that costs of reproduction exist in eusocial insect queens, but are latent, resulting in the positive fecundity-longevity relationship typically found in unmanipulated queens. We experimentally increased queens’ costs of reproduction by removing their eggs, which causes queens to increase their egg-laying rate. Treatment queens had significantly reduced longevity relative to control queens whose egg-laying rate was not increased. In addition, treatment and control queens differed in age-related gene expression based on mRNA-seq in both their overall expression profiles and the expression of ageing-related genes. Remarkably, this occurred principally with respect to relative age, not chronological age. These results provide the first simultaneously phenotypic and transcriptomic experimental evidence of costs of reproduction in queens of eusocial insects and suggest how the genetic pathways underpinning ageing might become remodelled during eusocial evolution.

## Introduction

The standard evolutionary theory of ageing (ETA) proposes that ageing occurs because individuals have limited resources to invest in maximising both fecundity and longevity, selection prioritises reproductive success over survival, and the strength of selection against ageing weakens with age^1–5^. Along with declining performance and increasing mortality with age (ageing), an outcome of these factors is a fecundity-longevity trade-off. Specifically, reproduction imposes costs by reducing fecundity and/or longevity later in life, leading to a negative fecundity-longevity relationship^4^.

The generality of this relationship has been challenged because, in theory, trade-offs between fecundity and longevity can become uncoupled and because, in some species, positive fecundity-longevity relationships occur^6–8^. For example, such relationships might occur within populations in which individuals vary in overall resource levels and well-resourced individuals invest in both high fecundity and high longevity. Nonetheless, in such populations a fecundity-longevity trade-off might still occur within individuals because no individual’s resources are limitless^9,10^.

Another potential exception is represented by eusocial insects such as eusocial Hymenoptera (ants, bees, and wasps with a worker caste) and termites. In these, reproductive phenotypes (queens or kings) are highly reproductive and comparatively long-lived, whereas workers are sterile or less reproductive and comparatively short-lived^11–14^. Furthermore, there appears to be a positive fecundity-longevity relationship within each caste, such that, in eusocial Hymenoptera, the most reproductive queens and workers (which can reproduce asexually in some species) live longer than the least reproductive queens and workers, respectively^15–26^. Therefore, it has been suggested that, in eusocial insects, queens (and reproductive workers) represent an exception to the usual negative fecundity-longevity relationship seen in other species, through not exhibiting costs of reproduction^22,24,27^. This apparent reversal of the conventional trade-off has been hypothesised to result from a remodelling of the conserved genetic and endocrine networks that regulate ageing and reproduction^13,28–33^. Recently, two such networks/gene sets have been proposed. One (the TI-J-LiFe network) contains the nutrient-sensitive target of rapamycin (TOR) and insulin/insulin-like signalling (IIS) pathways, which, in solitary insects, affect (via Juvenile Hormone signalling) ageing and reproduction through activation of immune and antioxidant pathways^33,34^. The other is a related enzymatic antioxidant gene set^35^. Both include age-related genes (genes that change expression with time) and ageing-related genes (genes in pathways that directly affect ageing) in several eusocial insects^33,35^.

In the eusocial bumblebee *Bombus terrestris,* the queens of which exhibit a positive relationship between lifetime reproductive success and longevity^16^, a recent study aimed to test experimentally for costs of reproduction in reproductive workers. It found that, as in other species, workers in unmanipulated colonies exhibited a positive fecundity-longevity relationship i.e. workers with more active ovaries lived longer^23^. However, this positive relationship was reversed (became negative) when randomly selected workers were experimentally manipulated to activate their ovaries. These results suggested that workers exhibit costs of reproduction but that workers choosing to reproduce in unmanipulated colonies are intrinsically high-quality individuals that can overcome such costs and achieve both high fecundity and longevity. In other words, costs of reproduction might be present but unexpressed (latent) in such individuals, implying condition-dependence of fecundity-longevity associations^23^. By extension, as queen-destined larvae receive nutrition of higher quality and/or in greater quantity^36^, such that adult queens are likely to be intrinsically well-resourced, high-quality individuals, the results suggested that queens also exhibit latent costs of reproduction. Hence, ageing and longevity patterns in queens could be explicable using ETA^23^.

Previous studies in the ant *Cardiocondyla obscurior* have found that age-related changes in gene expression in queens are opposite in direction to those described from female *Drosophila melanogaster*^28^ and have shown no effect on longevity of increased costs of reproduction in queens^24^. In the current study, therefore, we sought to test whether queens in eusocial insects experience costs of reproduction in an experiment that manipulated queens’ costs while profiling age-related gene expression changes in manipulated and control queens. Hence, we sought to test, simultaneously at phenotypic and transcriptomic levels, the extent to which eusocial insect queens represent a genuine exception to ETA. Specifically, using the *B. terrrestis* system, we aimed to discriminate between two alternative hypotheses. The first (H1) derives from ETA and states that a positive fecundity-longevity relationship in eusocial insect queens can arise because the costs of reproduction are present but latent. The second (H2) posits that eusocial insects represent a genuine exception to the trade-offs predicted by ETA because a remodelling of relevant genetic and endocrine networks has allowed queens to reproduce without costs^24,28,37^.

We manipulated queens’ costs of reproduction by experimentally removing eggs, which in *B. terrestris,* as in *C. obscurior*^24^, has been shown to increase queens’ fertility (realised fecundity, as measured by egg-laying rate)^38^. We allocated queens to two treatments. In Removal (R) queens, all eggs were removed from the colony, therefore inducing queens to increase their fertility and so experience greater costs of reproduction. In Control (C) queens, all eggs were removed from the colony and then replaced to control for disturbance (Figure 1a). We then measured the queen’s longevity in each colony. To control for potential effects of egg removal on workers’ aggression to queens^38^ and on queen activity levels, in both R and C colonies we also replaced aggressive workers with callow (newly-eclosed) workers^39^ and periodically measured queens’ locomotory activity and response to disturbance. To characterise changes in gene expression profiles with age brought about by the treatment, we sampled a subset of queens in both treatments at two time-points, the first when 10% queen mortality had occurred (time-point 1, TP1) and the second when 60% queen mortality had occurred (time-point 2, TP2) (Figure 1b). From each queen we extracted RNA from brain, fat body, and ovaries, since these tissues show gene expression changes with age in eusocial insects including *B. terrestris*^29,3,35,4^. We then used mRNA-seq to characterise changes in the gene expression profiles over time in both R and C queens and to compare the resulting *B. terrestris* gene lists with those from other studies in non-social and eusocial insects, including genes in the age-related TI-J-LiFe network and enzymatic antioxidant gene set^33,35^. H1 predicted that R queens should exhibit reduced longevity relative to C queens and a dissimilar pattern of change in the gene expression profile over time (as the egg-removal treatment should cause R queens to express their latent costs of reproduction). By contrast, H2 predicted that R and C queens should exhibit equal longevities and that their patterns of change in the gene expression profile over time should be similar (as costs of reproduction, latent or otherwise, are absent).

**Figure 1.**
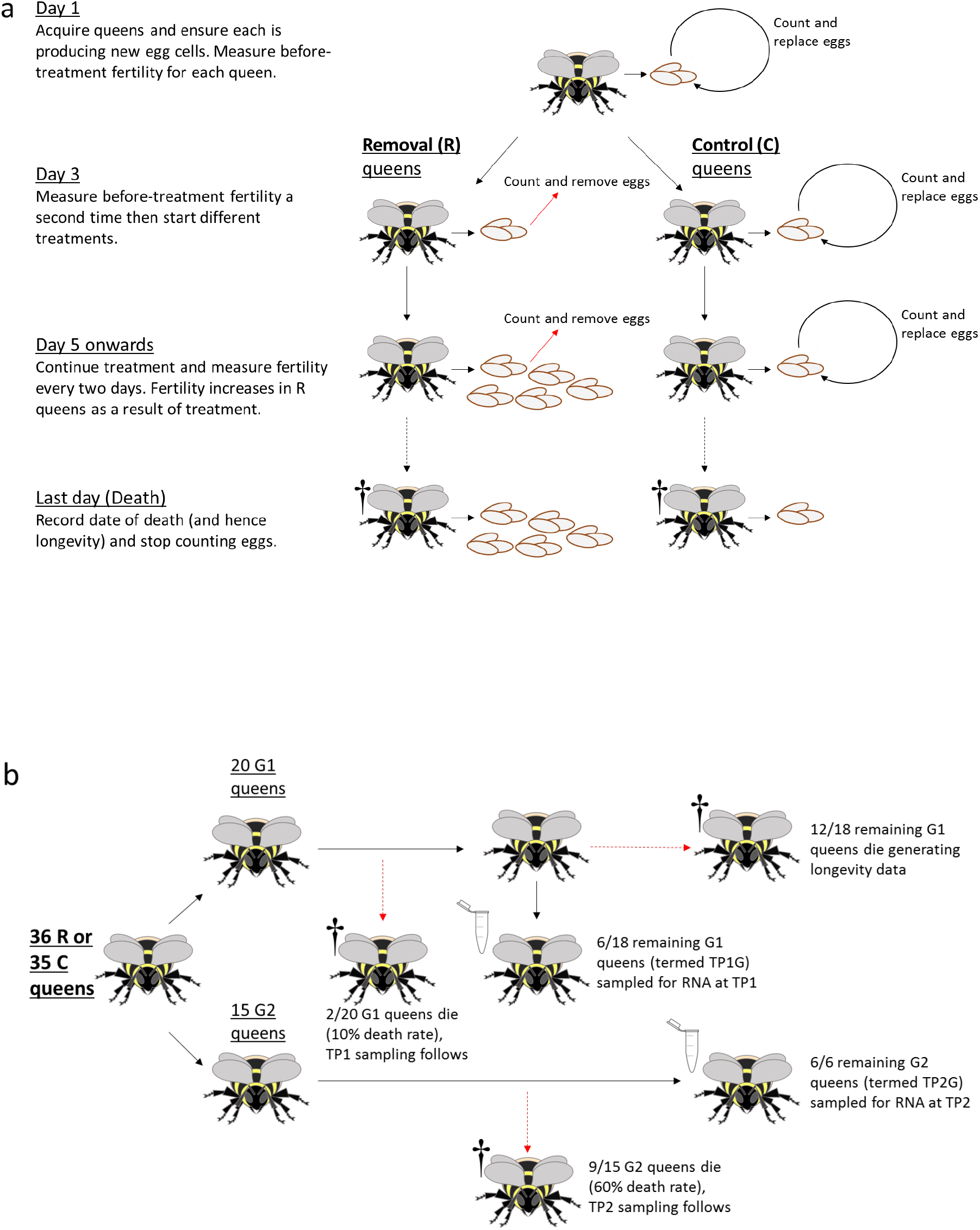
Outline of experimental design to test for costs of reproduction in *Bombus terrestris* queens. **a**, Experimental treatments: R: Removal queens: eggs removed and counted (straight red arrows); C: Control queens: eggs removed, counted, and replaced (circular black arrows). Day 1 was the first day that new egg-cells were observed in all colonies. **b**, RNA sample collection strategy. G1, G2: random subgroups of queens within treatments. TP1, TP2: time-points for sampling of queens for RNA extraction (R queens: TP1 on day 37, TP2 on day 85; C queens: TP1 on day 89, TP2 on day 134). TP1G, TP2G: subsets of queens sampled for RNA extraction at TP1 and TP2, respectively. One R queen (Q1) was not assigned to G1 or G2, and was used to provide life-history data only. One C queen (Q57 in the G1 subgroup) was censored and therefore not included in longevity analyses or sampled for RNA. Final sample sizes were: R:TP1G (N=6), R:TP2G (N=6), R:life-history (N= 2 + 9 + 12 + 1 = 24); C:TP1G (N=6), C:TP2G (N=6), C:life-history (N= 2 + 9 + 12 – 1 = 22). See Methods for full details.

## Results

### Queen fertility

There was no significant difference in baseline queen fertility (queen egg production before the treatment started) between R and C colonies (mean [95% CI] eggs per 48-hour period: R, 21.5 [18.8, 24.7]; C, 20.3 [17.7, 23.4]; negative binomial glmm: b = 0.057, SEb = 0.099, z = 0.578, p = 0.563; Figure 2a). However, there was a significant increase in queen fertility (queen egg production over days 5-25) across both groups (b = 0.893, SEb = 0.094, z = 9.485, p < 0.001) and this increase had a significant interaction with treatment (b = 0.650, SEb = 0.14, z = 4.766, p < 0.001), with R queens producing approximately twice as many eggs as C queens (mean [95% CI] eggs per 48-hour period: R, 52.5 [46.3, 59.6]; C, 25.9 [22.6, 29.7]) (Figures 2a, S1). R colonies also had higher colony fertility (overall colony egg production, including any worker-laid eggs) than C colonies (negative binomial glmm: b = 1.147, SEb = 0.078, z = 14.679, p < 0.001; Figures 2b, S2, S3; Table S1). Worker egg-laying (which occurred from day 26; Figures 2c, S4) accounted for only part of this difference (see Supplementary results for further details). Therefore, as intended, the R treatment increased queen egg-laying rate both before and after the start of worker egg-laying.

**Figure 2.**
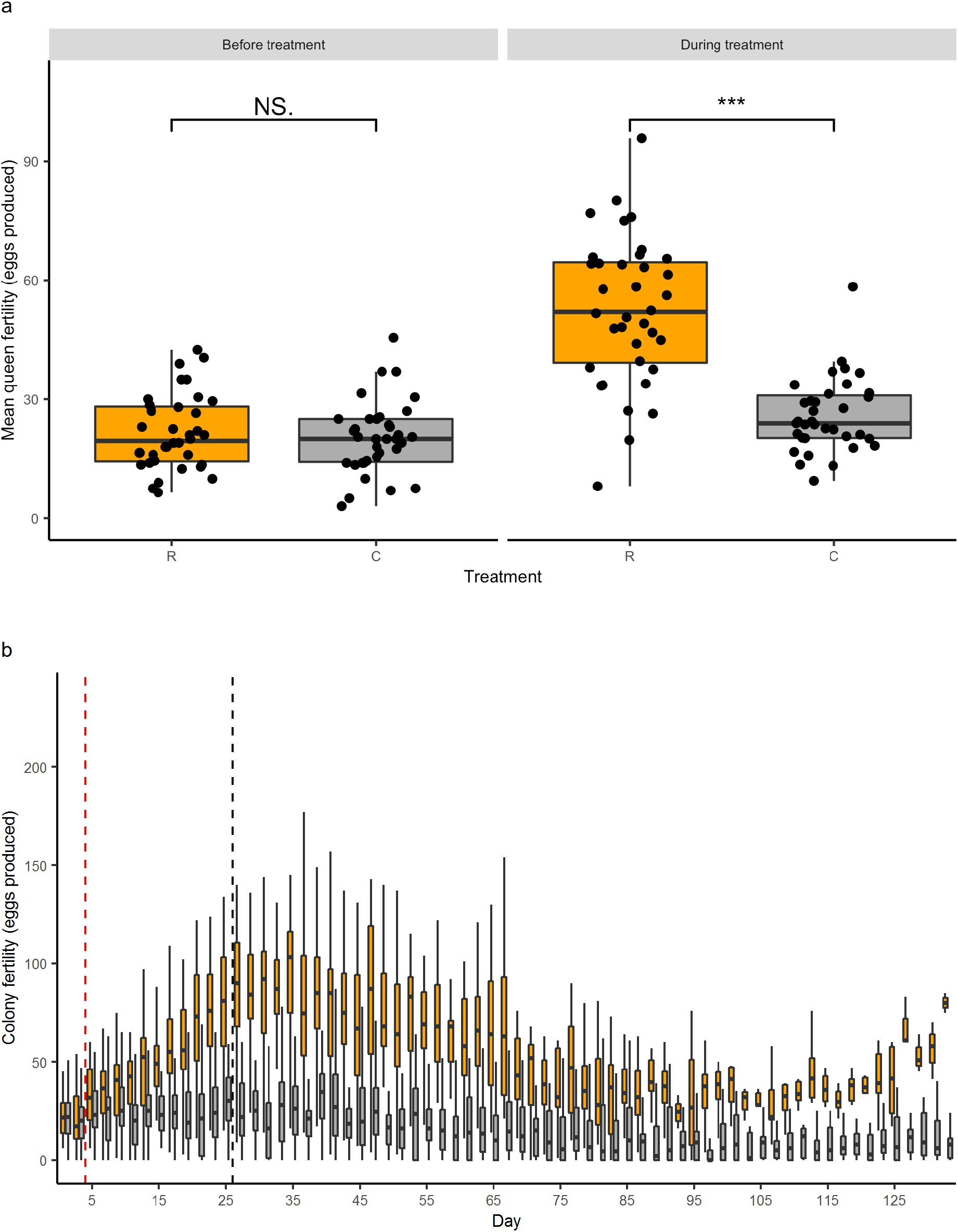
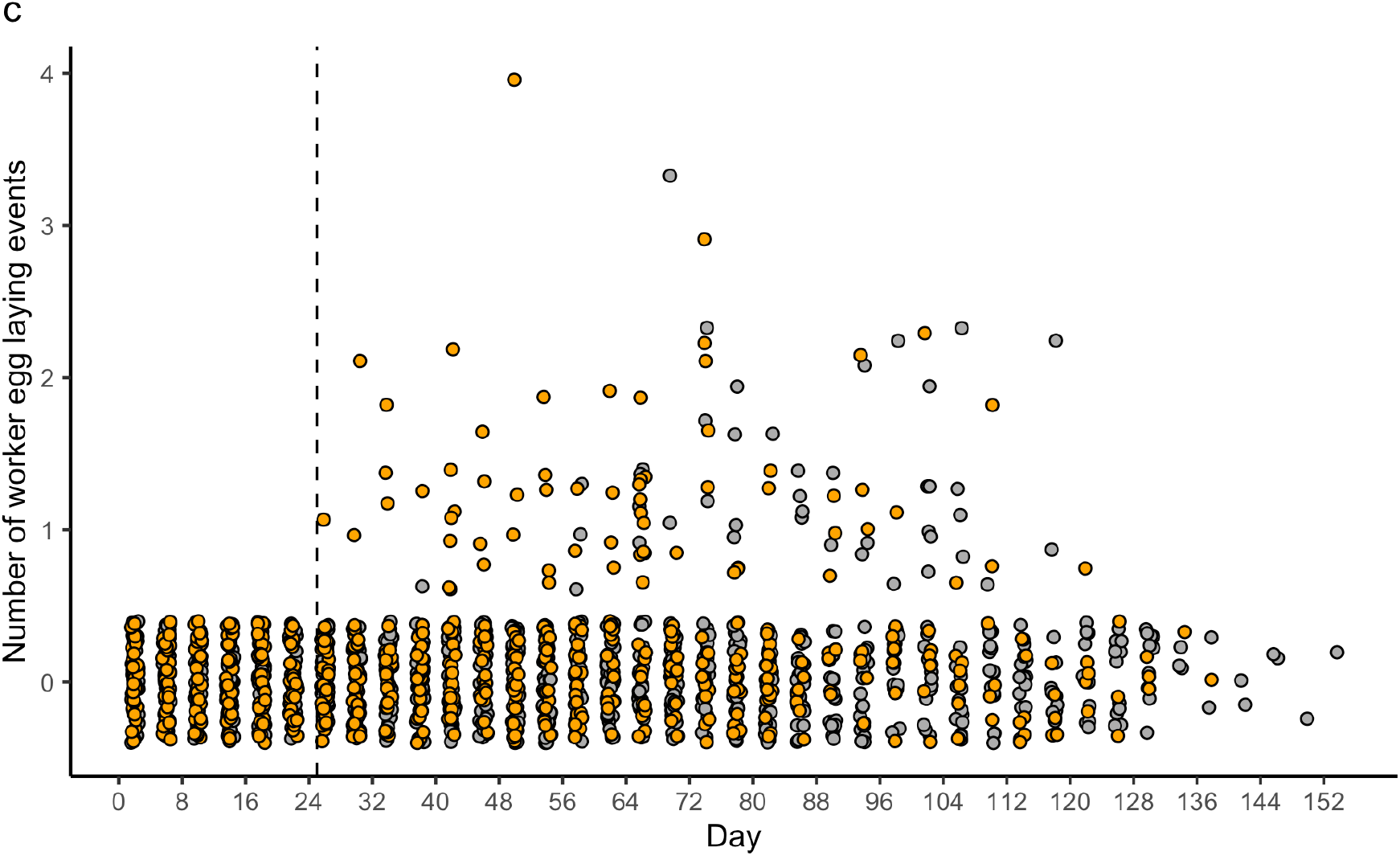
Fertility measures for R (eggs removed) and C (eggs removed and replaced) *Bombus terrestris* queens/colonies. **a**, Queen fertility (mean number of eggs produced per 48-hour period) for R (orange boxes; N = 36) and C (gray boxes; N = 35) colonies before treatment started (baseline fertility: days 1 and 3), and during treatment until the first worker egg-laying was observed (days 5-25 inclusive). NS, not significant; *** p < 0.001. Black circles represent means for individual queens. **b**, Colony fertility as a function of time for R (orange boxes; N = 36 on day 1 declining to N = 1 on day 134) and C (gray boxes; N = 35 on day 1 declining to N = 1 on day 148) colonies until day 134. Vertical red dotted line: day when manipulations started; vertical black dotted line: day 26 when worker egg-laying was first observed in any colony. Sample sizes on each day are in Table S28. Outliers are not shown. For **a** and **b**, black horizontal bars: median values for each treatment; boxes: interquartile ranges; whiskers: ranges up to 1.5 × the interquartile range. R colonies had significantly higher colony fertility than C colonies after treatment had started (see Results for details). **c**, Observed worker egg-laying as a function of time, recorded every four days during 10-minute observations in R (orange circles; N = 36 on day 1 declining to N = 1 on day 134) and C (gray circles; N = 35 on day 1 declining to N = 1 on day 148) colonies. Vertical black dotted line: day 26 when worker egg-laying was first observed in any colony. Points are offset around each integral value on the Y axis. From day 26 until day 74, R colonies had significantly higher numbers of worker egg laying events than C colonies (see Results for details).

### Queen longevity

R queens had significantly reduced longevity relative to C queens (median longevity in days [from day 1 of the experiment]: R, 73.6; C, 105.4; Cox’s proportional hazards analysis: hazards ratio = 0.224, z = −4.287, p < 0.001; Figures 3a, 3b). There was no significant difference in observed worker aggression (zero-inflated negative binomial glmm: b = 0.056, SEb = 0.307, z = 0.184, p = 0.854; Figure 3c) or filmed worker aggression (binomial glmm: b = 0.220, SEb = 1.18, z = 0.186, p = 0.853, Figure S5) between R and C colonies. In addition, observed worker aggression was significantly time-dependent for both treatments (zero-inflated negative binomial: glmm b = −0.057, SEb = 0.001, z = −5.782, p < 0.001), peaking at day 18 before declining throughout the rest of the experiment (Figure 3c), whereas mean queen longevity across both treatments was 89.4 days. Therefore, the reduction in queen longevity in the R treatment was not caused by greater worker aggression to queens. Nor was it caused by differences in queen activity levels or responsiveness between treatments, which were absent (Figures S6 - S8). Overall, these results support the prediction of H1 that queens in the R treatment, which experienced a greater cost of reproduction (from their higher egg-laying rates), should exhibit reduced longevity.

**Figure 3.**
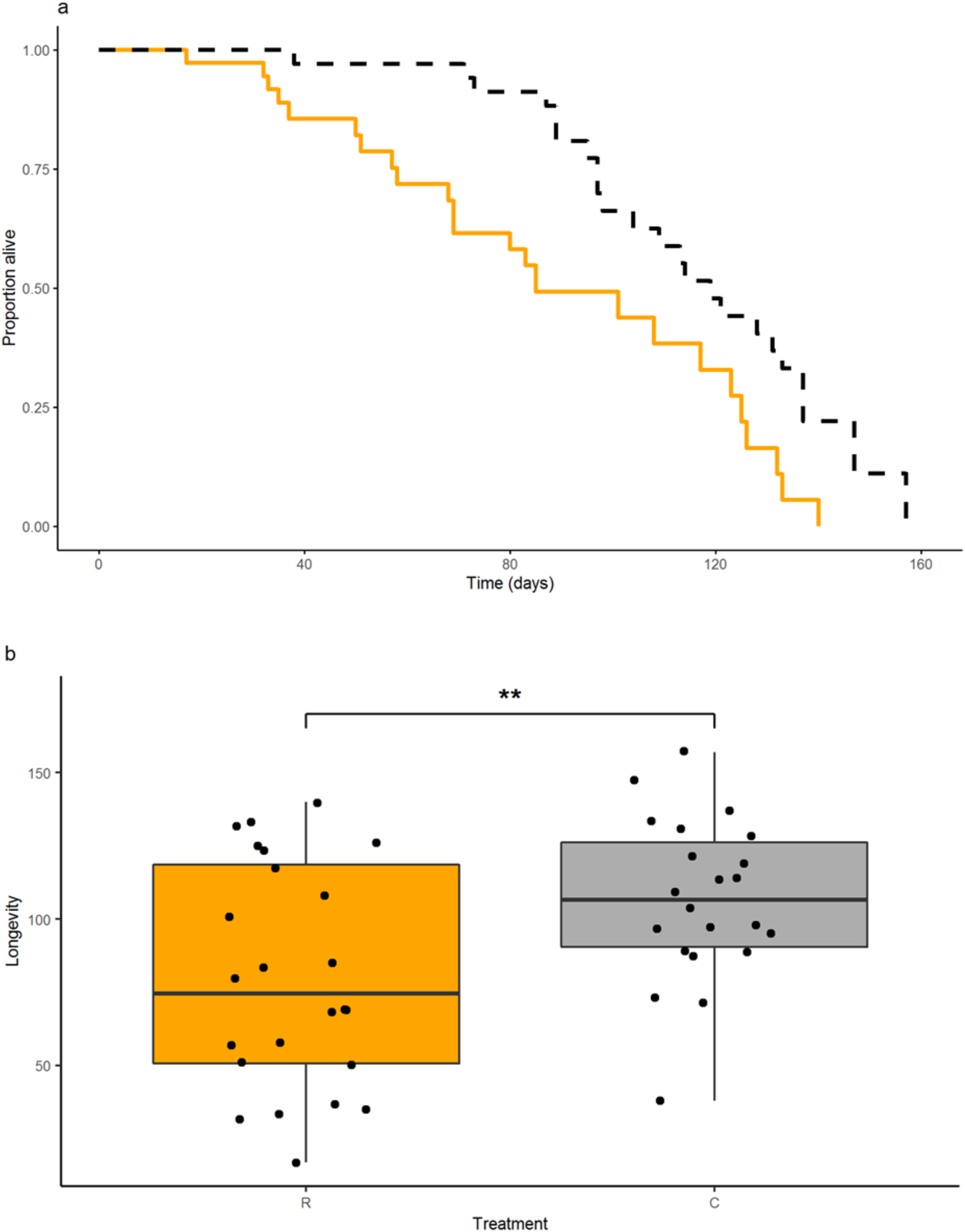
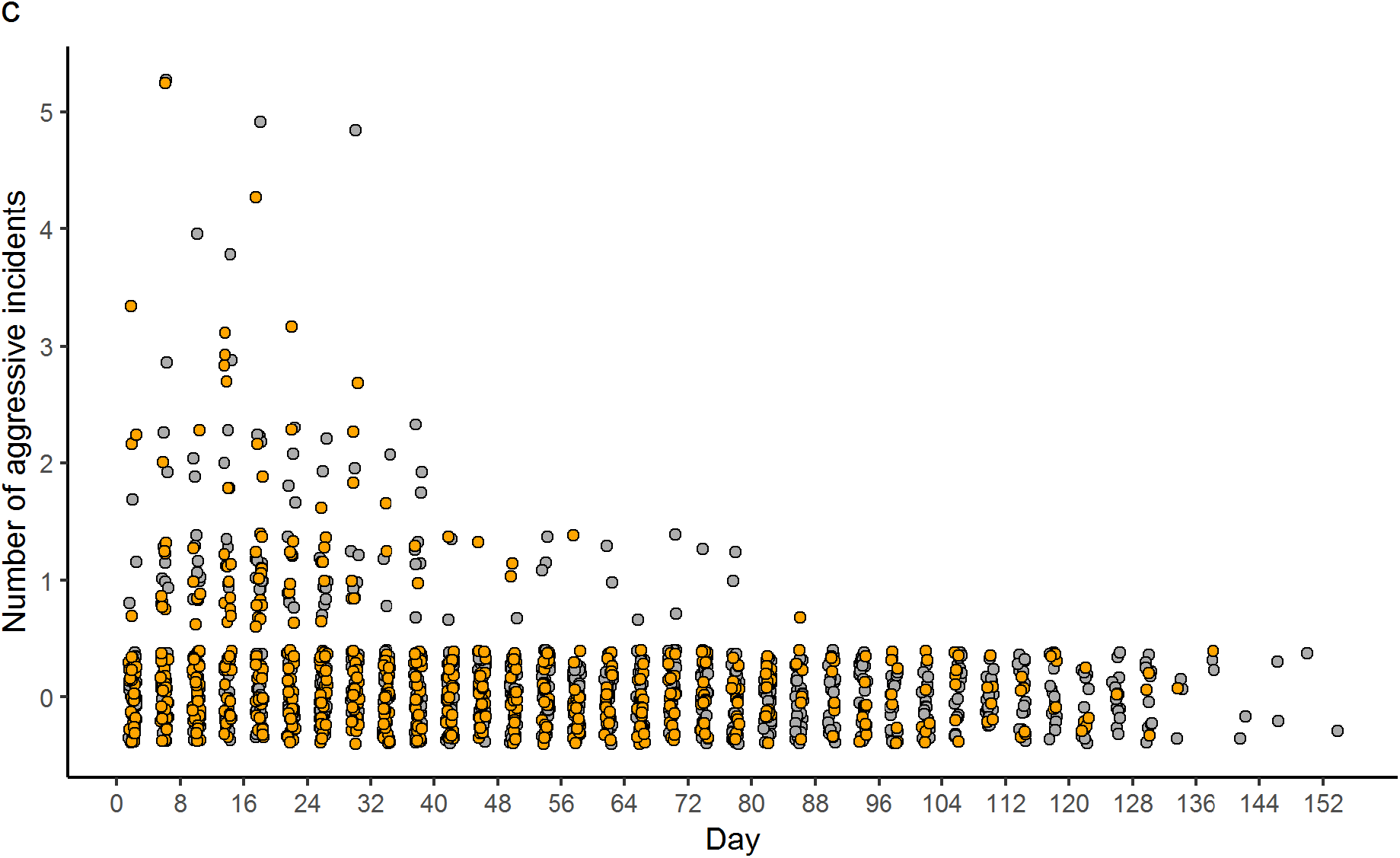
Queen longevity and worker aggression for R (eggs removed) and C (eggs removed and replaced) *Bombus terrestris* queens over the course of the experiment. **a**, Survivorship curves for R (solid orange line, N = 24) and C (dotted black line, N = 22 queens); **b**, Boxplots showing longevity (days) for R (orange boxes, N = 24) and C (gray boxes, N = 22 queens). Black horizontal bars: median queen longevity; boxes: interquartile ranges; whiskers: ranges up to 1.5 × the interquartile range. ** p < 0.01. For figures **a**, and **b**, R queens had significantly reduced longevity relative to C queens (see Results for details); **c**, Observed worker-to-queen aggression as a function of time (recorded every four days during 10-minute observations) in R (orange circles, N = 36 on day 1 and N = 1 on day 134) and C (gray circles, N = 35 on day 1 and N = 1 on day 148) colonies. Sample sizes on each day are in Table S28. Points are offset around each integral value on the Y axis. There was no significant difference in observed worker-to-queen aggression between R and C colonies (see Results for details).

### Age-related gene expression

The mRNA-seq gene expression profiles (Figures S9 - S11; Tables S2 – S11; see Supplementary results for further details) showed that, in each tissue (brain, fat body, ovaries), R queens exhibited far less differential gene expression between TP1 and TP2 than C queens (Figures 4, 5, S9 - S11). To determine more exactly whether gene expression profile changes over time (from TP1 to TP2) between R and C queens differed (H1) or not (H2), we compared lists of differentially expressed genes (DEGs) across R and C treatments within each tissue (Tables S4 – S6). Overall, 3/6 comparisons of R and C DEGs showed no significant overlap and 3/6 showed significant overlap (Figure 4; Tables S12, S13). The significant overlaps were: (a) in brain, 9/30 of the DEGs that had significantly higher expression in TP2G than TP1G (‘up-regulated’ DEGs) in R were also up-regulated in C (Fisher’s exact test, p = 5.33 x 10^-6^; Figure 4a; Table S12); (b) in brain, 4/7 of the DEGs that had significantly higher expression in TP1G than TP2G (‘down-regulated’ DEGs) in R were also down-regulated in C (Fisher’s exact test, p = 2.72 x 10^-5^; Figure 4b; Table S12); and (c) in fat body, 77/412 of the DEGs down-regulated in R were also down-regulated in C (Fisher’s exact test, p = 2.16 x 10^-13^; Figure 4d; Table S12). However, levels of overlap were relatively low and there were at least double the number of DEGs (up- or down-regulated) in C compared to R queens in each tissue (Figure 4). Gene ontology (GO) enrichment analysis showed that the biological functions of age-related DEGs (in brain and fat body) also differed between R and C queens (Table S7; see *Gene Ontology (GO) enrichment analysis,* Supplementary results, for further details). Hence, overall, R and C queens exhibited dissimilar patterns of change in both total gene expression profiles and DEG functions with their relative age (age measured in terms of percentage mortality).

**Figure 4.**
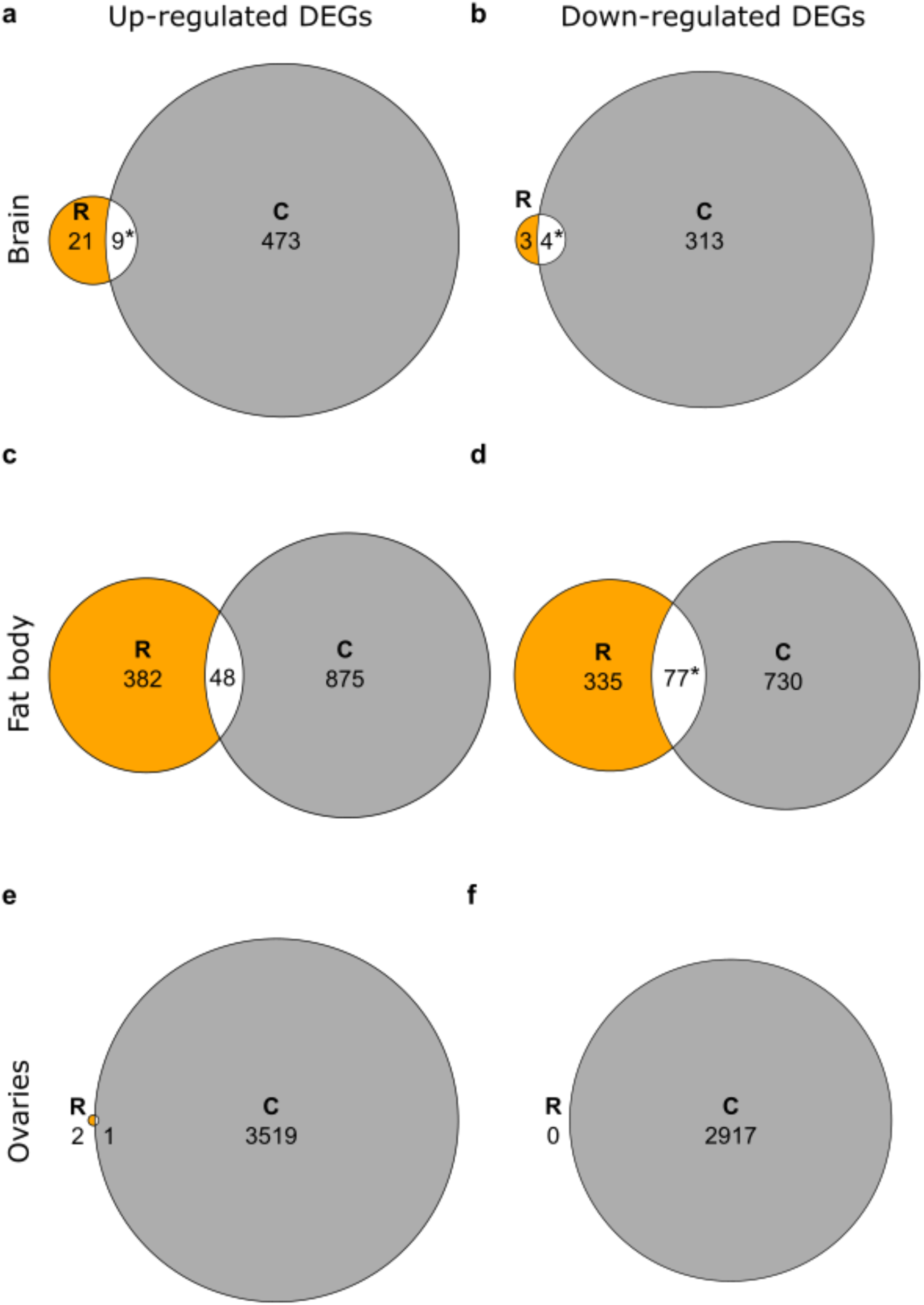
Comparison of changes in gene expression profiles with time in three tissues between R (eggs removed) and C (eggs removed and replaced) *Bombus terrestris* queens as determined by mRNA-seq. Euler diagrams of overlaps between differentially expressed genes (DEGs), i.e. genes differentially expressed between the two time-points TP1 and TP2 and shared between R queens (orange circles) and C queens (gray circles) for: **a**, **b**, brain; **c**, **d**, fat body; and **e**, **f**, ovaries. (In panel **e**, because their low values mean there is a lack of space, numbers of DEGs for R queens and shared between R and C queens are shown adjacent to the relevant area.) Asterisks (*) denote significant overlap in DEGs (Fisher’s exact test, *p* < 0.05 after Bonferroni correction). Up-regulated DEGs: DEGs significantly more expressed in TP2G than TP1G, i.e. that increase expression with queen age; down-regulated DEGs: DEGs significantly more expressed in TP1G than TP2G, i.e. that decrease expression with queen age. Results of statistical tests are in Table S12 and identities of overlapping genes are in Table S13.

**Figure 5.**
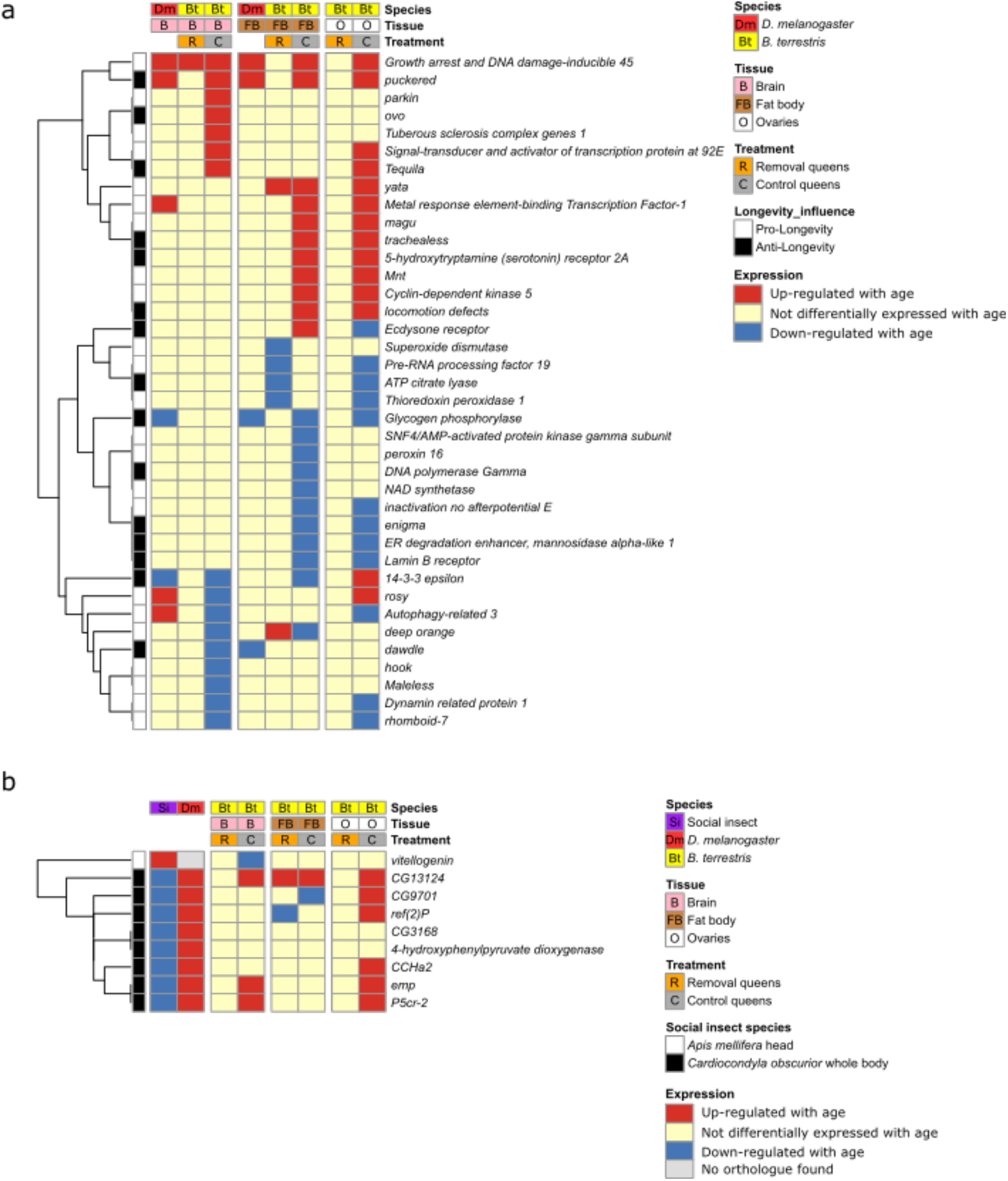
Gene expression patterns compared between age-related genes in *Bombus terrestris* queens (current study) and age and/or ageing-related genes in other insect species. **a**, Comparison of *B. terrestris* with *Drosophila melanogaster* for genes in the *D. melanogaster* GenAge database of pro- and anti-longevity genes. (As the GenAge database does not specifiy patterns of age-related gene expression, data on age-related gene expression from tissue-specific *D. melanogaster* studies (brain = Pacifico et al.^42^, fat body = Chen et al.^41^ are included in the figure for comparison. Data on expression in *B. terrestris* ovaries from the current study are shown for illustration, although no comparable data were available on *D. melanogaster* age-related gene expression in ovaries.) Each row represents an individual gene showing age-related differential expression in brain or fat body of *B. terrestris* (current study) and having a single-copy orthologue present in the *D. melanogaster* GenAge database as determined by Orthofinder (N=38 unique genes from the 46 overlapping genes in Tables S16, S17). Each column denotes whether a gene shows age-related differential expression in a given data set (brain: *D. melanogaster* = Pacifico et al.^42^, *B. terrestris* = current study; fat body: *D. melanogaster* = Chen et al.^41^, *B. terrestris* = current study; ovaries: *B. terrestris* = current study). **b**, Comparison of *B. terrestris* with *Apis mellifera, Cardiocondyla obscurior* and *D. melanogaster* genes having opposite patterns of age-related expression change in *C. obscurior*. Each row represents an individual gene from *A. mellifera* or *C. obscurior* whose expression in queens is age-related and has a single-copy orthologue in *B. terrestris*, as determined by Orthofinder (N = 9 genes). Each column denotes whether a gene is differentially expressed in a given data set (*A. mellifera* = Corona et al.^66^; *C. obscurior* = von Wyschetzki et al.^28^; *D. melanogaster =* Pletcher et al.^68^ and Doroszuk et al.^61^, as reported by von Wyschetzki et al.^28^; *B. terrestris* = current study). In both **a** and **b**, vertical breaks separate the three tissues (brain, fat body, and ovaries) and the dendograms at left group genes that cluster together according to their gene expression patterns.

R and C queens might have differed in their total age-related gene expression profiles because sampling queens at identical time-points with respect to relative age led to the periods between the sampling time-points differing between the treatments with respect to chronological age (absolute age, here measured in days from day 1) (Figure 1). We therefore determined whether *B. terrestris* R and C queens differed from one another as regards their expression patterns for age- and/or ageing-related genes identified in previous studies of solitary *(D. melanogaster)* and eusocial (the honeybee *Apis mellifera* and the ant *C. obscurior)* insect species. For *D. melanogaster,* no comparisons in either R or C queens showed significant overlap between orthologues of DEGs and (i) comparable tissue-specific mRNA-seq *D. melanogaster* studies^41,42^ (0/8 comparisons; Tables S14, S15) or (ii) the *D. melanogaster* GenAge database^43^ (0/6 comparisons; Tables S16, S17) (Figure 5a). Therefore, in these comparisons, *B. terrestris* queens (across both treatments) and *D. melanogaster* females did not exhibit a shared set of age- and/or ageing-related genes, rendering the comparisons uninformative as regards discriminating H1 and H2. For the *A. mellifera* age-related gene *vitellogenin,* R and C queens differed in that R queens showed no age-related differential expression and C queens showed age-related differential expression in the opposite direction compared with *A. mellifera* queens (Figure 5b). In comparisons with a set of eight age-related genes (with single-copy orthologues in *B. terrestris*) in the ant *C. obscurior* having opposite expression patterns in *D. melanogaster*^28^, *B. terrestris* R queens showed no significant overlaps (in brain, fat body, or ovaries) with either species (Figure 5b; Table S18). However, *B. terrestris* C queens showed significant overlap in brain (but not in fat body or ovaries) with these same eight genes in *D. melanogaster* but not *C. obscurior* (Figure 5b; Table S18).

In comparisons of DEGs with the TI-J-LiFe network^33^, R queens did not significantly overlap in any of the three tissues, whereas C queens showed significant overlaps in the brain (for 5/6 comparisons) but not fat body or ovaries (Figure S12; Tables S19, S20). For the enzymatic antioxidant gene set^35^, R queens showed significant overlap in fat body (though for only one comparison, i.e. when the top 300 genes from each list were compared) but not in brain or ovaries, whereas C queens showed no significant overlaps in any of the three tissues (Figure S13; Tables S21, S22).

To investigate further whether the age-related gene expression patterns observed between R and C queens were affected by sampling at time-points reflecting relative rather than chronological age, we determined the number of genes that were significantly differentially expressed between R:TP1 (day 37) and C:TP1 (day 89), between R:TP2 (day 85) and C:TP1 (day 89), and between R:TP2 (day 85) and C:TP2 (day 134) (Figures 6, S14; Tables S23 - S25). Across the three tissues, between R:TP1 and C:TP1, there were moderate numbers of DEGs (15-1077 up-regulated, 6-986 down-regulated); between R:TP2 and C:TP1, there were few DEGs (0 up-regulated, 2-5 down-regulated); and between R:TP2 and C:TP2, there were large numbers of DEGs (728-3433 up-regulated, 645-2611 down-regulated) (Figures 6, S14; Tables S23 - S25). In addition, there were significant and high percentage overlaps in DEGs for which the time-period (the span between two time-points) also overlapped (e.g. R:TP1 to C:TP1 versus R:TP1 to R:TP2; Tables S26, S27; see Supplementary results for details). The 7 DEGs between R:TP2 and C:TP1 had no single-copy orthologues in *D. melanogaster*, and therefore it was not possible to ascertain if they were represented in either the TI-J-LiFe network or the enzymatic antioxidant gene set (Tables S23, S25).

**Figure 6.**
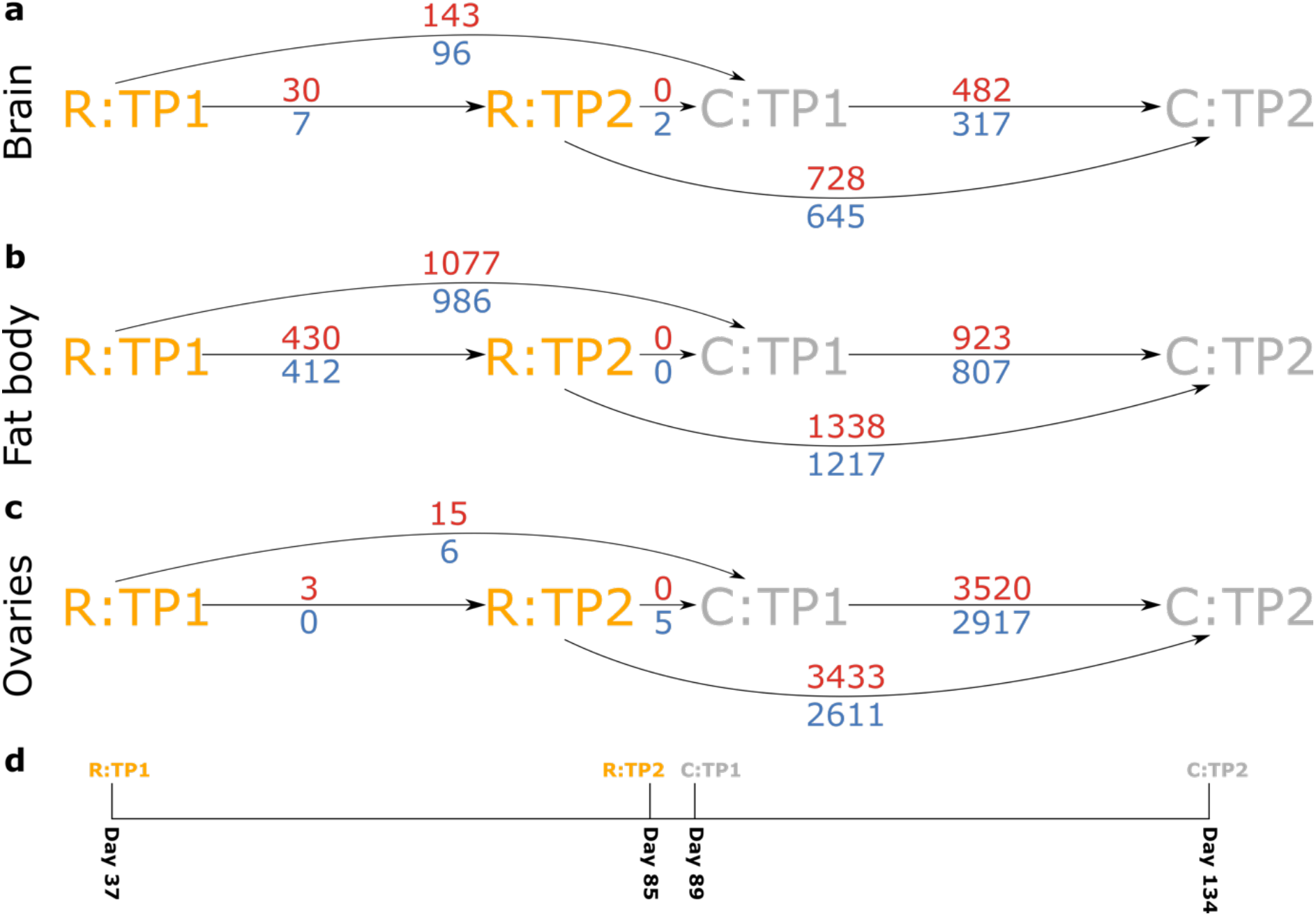
Comparisons in gene expression profiles between chronological ages in three tissues of R (eggs removed) and C (eggs removed and replaced) *Bombus terrestris* queens as determined by mRNA-seq. Arrows and associated numbers denote differentially expressed genes (DEGs) between two conditions (increasing in chronological age from left to right) for **a,** brain; **b,** fat body; and **c,** ovaries. R conditions are represented in orange and C conditions are represented in gray. Red numbers, above arrows: numbers of genes up-regulated (more expressed with chronological age) between two conditions linked by an arrow; blue numbers, below arrows: numbers of genes down-regulated (less expressed with chronological age) between two conditions linked by an arrow. **d,** Scale displaying the day of the experiment on which each condition was sampled. TP1: time-point 1, TP2: time-point 2.

As the gene expression differences between R:TP2 and C:TP1 were overall much smaller than for the other comparisons, and as queens sampled at these time-points had similar chronological ages but different treatments and relative ages (R:TP2: 85 days, 60% mortality; C:TP1: 89 days, 10% mortality), these and the previous results (Figure 4) imply that age-related changes in gene expression profile caused by the experimental treatment occurred predominantly with respect to queens’ relative age and not their chronological age. However, although R:TP2 and C:TP1 exhibited similar expression profiles, the results were inconsistent with there being a common gene expression trajectory for all queens irrespective of treatment. This can be concluded, despite the high percentage overlaps, because, in brain and fat body (though not ovaries), there were many more DEGs isolated between R:TP1 and C:TP1 than between R:TP1 and R:TP2, and there were many more DEGs isolated between R:TP2 and C:TP2 than between C:TP1 and C:TP2 (Figures 6, S14).

Overall, therefore, the mRNA-seq analyses supported the prediction of H1 that R and C queens should exhibit a dissimilar pattern of change in the gene expression profile over time but also suggested a degree of invariance of gene expression profile change with chronological age.

## Discussion

We experimentally manipulated queens of the eusocial bumblebee *B. terrestris* by removing their eggs, causing them to double their egg-laying rate. This increase in the costs of reproduction led to a significant decrease in queen longevity, which fell by 30% relative to that of control queens (to 73.6 days from 105.4 days, as measured from the experiment’s start day). The decreased longevity of treatment queens was not caused by increased worker aggression, queen activity, or queen responsiveness. In addition, treatment queens differed from control queens in the pattern of change in their gene expression profiles with relative age, as regards both overall gene expression and the expression of ageing-related genes. Our results therefore support the hypothesis (H1) that a positive fecundity-longevity relationship in eusocial insect queens can arise because costs of reproduction are present but latent, with the high individual quality inherent in the queen phenotype allowing queens in normal conditions to overcome such costs and exhibit both high fecundity and high longevity. This explanation is consistent with ETA (see Introduction) and our results therefore imply that eusocial insects are not universally an exception to the predicted fecundity-longevity trade-off found in most non-social species. However, our results showing that age-related changes in gene expression profile exhibited a degree of invariance with chronological age also suggest, consistent with H2, that some remodelling of genetic and endocrine networks underpinning ageing has occurred in *B. terrestris*, as we discuss further below.

The difference between the results of Schrempf et al.^24^ for *C. obscurior* queens, which showed no effect on longevity of increased costs of reproduction, and the current results for *B. terrestris* queens, conceivably arises from differences in the colony cycle and/or level of eusocial complexity. *C. obscurior*, like all ants and termites, has a perennial colony cycle in which queens potentially live many years^44^. By contrast, *B. terrestris* like all temperate eusocial bees and wasps except for *Apis* spp., has an annual colony cycle in which queens, though still longer lived than workers, live for approximately one year^45,46^. Associated with this difference, perennial species exhibit a more ‘advanced’ degree of eusociality relative to the ‘intermediate’ or ‘primitive’ eusociality of annual species^47–50^. It is possible, therefore, that avoidance of costs of reproduction, accompanied by a complete remodelling of relevant genetic and endocrine pathways, occurs in queens of perennial, advanced eusocial insects but not those of annual, less advanced ones^13^.

Gene expression changes with relative age in R and C queens differed in exhibiting: (i) dissimilar overall profile changes; (ii) unalike GO terms; (iii) different patterns of expression change in *vitellogenin;* (iv) overlaps in C but not R queens with the eight *D. melanogaster* genes having opposite patterns of age-related expression change in *C. obscurior;* and (v) differential overlaps with ageing-related genes from the TI-J-LiFe network and enzymatic antioxidant gene set. These results supported H1’s prediction that the treatment would cause differences in age-related gene expression profiles between R and C queens. Given that R and C queens differed in the costs of reproduction and longevity, they also supported the suggestion that modulation of genes in the TI-J-LiFe network and the enzymatic antioxidant gene set underpins fecundity-longevity relationships and mechanisms of ageing in eusocial insects^33,35^.

Intriguingly, there was a close similarity in gene expression profiles across treatments between queens of differing relative ages (R:TP2 queens, 60% mortality vs C:TP1 queens, 10% mortality) but similar chronological ages (85 vs 89 days old, respectively). Although there appeared not to be a common gene expression trajectory for both R and C queens, queens of almost identical chronological age showed similar gene expression profiles despite their different costs of reproduction. These results imply that age-related changes in gene expression profile caused by the treatment occurred predominantly with respect to queens’ relative age and not their chronological age. Such invariance with chronological age appears absent in solitary insect species^51,52^. Therefore, these results suggest that while *B. terrestris* queens can indeed exhibit costs of reproduction (as H1 predicted), they may have undergone a partial remodelling of genetic pathways underpinning ageing such that the overall profile of age-related gene expression depends more on chronological age than relative age. This could be associated with the intermediate eusocial level of *B. terrestris* and similar taxa, and conceivably represents a step in the route to the complete remodelling hypothesised to have occurred in more advanced eusocial species^24^.

In conclusion, experimental life-history and gene expression profiling data from *B. terrestris* support the occurrence of costs of reproduction in annual eusocial insects of intermediate social complexity and suggest that eusocial insect queens may exhibit condition-dependent positive fecundity-longevity associations. They also suggest that some degree of remodelling of the genetic and endocrine networks underpinning ageing has occurred in intermediately eusocial species. Therefore, as others have suggested^53,54^, understanding costs of reproduction and their genetic underpinnings remains central to explaining the evolutionary trajectory of the effects of eusociality on ageing and life history.

## Methods

### Colony maintenance

We obtained 75 young *B. t. audax* colonies (mean [SD] number of workers = 9.1 [3.9]) from Biobest Group NV (Westerlo, Belgium) on 28 March 2019. The queens of these colonies had been placed into hibernation on 22 October 2018 and their hibernation had ended on 18 February 2019 (Annette Van Oystaeyen, Biobest Group NV, personal communication). On receipt we transferred all colonies to wooden nest-boxes (17 × 27.5 × 16 cm) with clear Perspex lids. We kept colonies at 28°C and 60% RH under constant red light and provided them with *ad libitum* sugar solution and pollen throughout the experiment. We set aside four colonies as ‘source colonies’ because either their queen had died on arrival (two colonies) or they contained more than 20 workers (two colonies). The remaining 71 ‘experimental colonies’ contained a living queen and 3 – 17 workers (mean 8.4 workers per colony) on receipt. We used the source colonies, supplemented by four additional source colonies obtained on 3 July 2019, to provide callow workers and brood to the 71 experimental colonies throughout the experiment.

To assign the experimental colonies to each treatment and to control for the effect of initial colony size on queen fertility and longevity, we paired each colony with the colony closest to it in size, producing 35 colony pairs, and then randomly selected one colony from each pair for each treatment. The single unpaired colony remaining was randomly assigned to the R treatment. Hence this procedure created 36 R (Removal) colonies and 35 C (Control) colonies.

For the remainder of the experiment, to control colony size and facilitate manipulations, we maintained the number of workers within each experimental colony in both R and C treatments at 20 workers, which lies within the range of sizes found in field *B. terrestris* colonies (mean [range] number of workers = 35.6 [11-75])^55^. To achieve a constant worker number, we marked each worker initially present with white Tipp-Ex (Bic, Île-de-France, France) applied to its thorax. We then continued to mark and count workers as they eclosed every day until the colony acquired 20 workers (removing excess workers, identified as any workers without marks, after this point). As some excess workers were callows (identifiable from their less pigmented coats), we used them together with callow workers from the source colonies to bring the number of workers in colonies with fewer than 20 workers to the required level (*B. terrestris* colonies will accept non-nestmate callow but not adult workers^56^. Once a colony had 20 workers, we continued to remove excess workers or to add callow workers to maintain worker number at 20 per colony. In addition, we removed all dead workers and, as they eclosed, all new (virgin) queens and males. In each case, to assess any effect of worker removals and deaths on queen longevity, we recorded the numbers of excess and dead workers removed from each colony and callow workers added to each colony. We found no significant effect of treatment on numbers of dead workers removed (mean [SD] workers per colony: R, 8.5 [7.8]; C, 11.5 [8.4]; negative binomial glmm: b = −0.146, SEb = 0.187, z = −0.780, p = 0.435) or callow workers added (mean [SD] workers per colony: R, 8.8 [7.1]; C, 11.3 [8.2]; negative binomial glmm: b = −0.226, SEb = 0.200, z = −1.128, p = 0.259). However, as expected, there were significantly fewer excess workers removed from R than from C colonies (mean [SD] workers per colony: R, 39.1 [19.1]; C, 67.9 [30.7]; negative binomial glmm: b = 0.373, SEb = 0.126, z = 2.971, p = 0.003).

### Experimental manipulation and colony fertility

To ensure that all experimental colonies could produce eggs, we defined day 1 of the experiment as the first day on which new egg-cells were present in every colony across all colonies (1 April 2019). To monitor new egg-cell production, we used a marker pen to map, on a clear acetate sheet fastened to the Perspex lid of the nest-box, the positions of all egg-cells for each colony daily. To measure colony fertility (egg-laying rate from day 1 until the last queen death on day 158), we collected data from each experimental colony on the numbers of egg-cells and the number of eggs within them every two days (‘count days’). These included: 1) two count days (days 1 and 3) before the R and C treatments were started (on day 3); and 2) every count day until each queen’s death. The day 1 and 3 counts were conducted to measure baseline queen fertility before treatments were begun. For these counts, we carefully removed and opened all new egg-cells in each colony, counted the eggs in each cell, resealed the cells, and placed them back into their previous positions in each colony. From day 3, we implemented the two treatments as follows: 1) R: we removed and opened each new egg-cell, counted the eggs inside, removed the eggs, resealed the empty cell, and placed it back into its previous position in the colony; 2) C: we removed and opened each new egg-cell, counted the eggs inside, and, leaving the eggs in place, resealed the cell and placed it back into its previous position in each colony. We used these count data to produce two separate indices of queen fertility for each colony: 1) baseline queen fertility, i.e. the mean of the egg counts across each count day before experimental manipulations had begun (days 1 and 3); and 2) during-treatment queen fertility, i.e. the mean of the egg counts across each count day after experimental manipulations had begun up to the point when worker egg-laying was first observed (days 5-25). Some *B. terrestris* workers lay unfertilised, male eggs at a characteristic point in the colony cycle, the ‘competition point’^57^, and the first of these in any colony was observed on day 26; hence measures of queen/colony fertility were separated into those taken before and after day 26, as the latter would have included a mixture of queen- and worker-produced eggs. As we observed no worker egg-laying before day 26, and as removal of queen-laid eggs does not affect the timing of the competition point^38^, we counted all eggs laid before day 26 as queen-laid eggs (i.e. queen fertility).

### Queen longevity

To measure queen longevity, we checked each queen daily. Once a queen had died, we recorded her date of death, removed her, and froze her and her colony at −20°C. Therefore, queen longevity was calculated as the number of days between day 1 and the queen’s date of death. One C queen (Q57) escaped during the count process and was therefore excluded from further analysis.

### Observed queen activity

As queen longevity was potentially affected by queen activity level, every four days (‘observation days’, which occurred on days without egg counts/manipulations), we recorded whether each queen was active (walking/running or engaged in egg-laying/aggression behaviours) or inactive (not engaged in these behaviours) during a brief (i.e. of a few seconds) observation period. These data were used to calculate ‘observed queen activity’ (proportion of queens that were classified as active, in each treatment, per observation day). After the observation period we physically carried the nest-box to a monitoring station and then recorded the queen’s response (previously active queen increases her movement speed and previously inactive queen becomes active, as defined above) or non-response (none of these events occurs). These data were used to calculate ‘response to disturbance’ (proportion of queens that responded to disturbance relative to those that did not respond to disturbance, in each treatment, per observation day).

### Observed worker aggression and observed worker egg laying

In *B. terrestris*, removing queen eggs increases worker-to-queen aggression^38^ and worker egg-laying is associated with worker-to-queen and worker-to-worker aggression^58^. We therefore sought to equalise worker aggression across R and C treatments by removing egg-laying and/or aggressive workers from all colonies^39^. For this, every four days, following each response to disturbance test (above), we observed the workers in each colony for 10 minutes. During this observation period we recorded ‘observed worker egg-laying’ (defined as the number of worker egg-laying events observed in this period per colony per observation day) and ‘observed worker aggression’ (defined as the number of aggressive incidents directed by workers^59^ towards the queen observed in this period per colony per observation day). We recorded worker-to-queen but not worker-to-worker aggression as only worker-to-queen aggression was expected to have effects on queen longevity. Queen-to-worker aggression was not observed at any stage during the experiment. We also removed any egg-laying worker or aggressive (towards the queen) worker that was observed during the observation period and replaced it with a marked callow worker from a source colony or another experimental colony. Time spent removing and replacing workers was not included in the 10-minute observation period. Following these manipulations, the number of workers removed due to egg-laying was significantly higher in R than in C colonies, although low in both cases (mean [SD] workers per colony: R, 2.3 [2.5]; C, 1.5 [1.2]; negative binomial glmm: b = −0.991, SEb = 0.219, z = −4.531, p < 0.001); the number of workers removed due to aggression did not differ significantly between treatments and was also low (mean [SD] workers per colony: R, 1.6 [1.7]; C, 2.1 [2.6]; negative binomial glmm: b = −0.135, SEb = 0.291, z = −0.462, p = 0.644).

### Digital filming

To supplement the data on queen and worker behaviour obtained from the direct observations, we also filmed each colony using Sony CDR-CX190 digital camcorders (Sony, Tokyo, Japan) under white light for one hour on two days during the experiment. These two days (days 48 and 99) were chosen to coincide with TP1 and TP2 in the C queens (Figure 1b; see also *Collection of queens for mRNA-seq* below). The films were viewed during playback and the following metrics were quantified using the behavioural software package *BORIS* ^60^: ‘filmed queen activity’ (the amount of time a queen was active during filming, per colony, per film period; activity defined as for observed queen activity); ‘filmed worker egg laying’ (the number of worker egg-laying events recorded during filming, per colony, per film period); and ‘filmed worker aggression’ (number of worker aggressive incidents that were caught on film, per colony, per film period; aggressive incidents defined as for observed worker aggression). All film data were collected blindly with respect to colony treatment. We filmed all colonies whose queens were still alive at the time of each film period. These comprised 25 R colonies and 33 C colonies in the first film period (day 48) and 8 R colonies and 20 C colonies in the second film period (day 99). We analysed film from all R colonies (N=8) filmed in both periods and a set of C colonies (N=9) randomly selected from the 20 C colonies filmed in both film periods.

### Brood transfer between colonies

Because R colonies produced no larvae (as all their eggs were removed), we equalised the number of third/fourth-instar larvae and pupae across R and C colonies. We did this by first removing some of the larvae and pupae from the C and source colonies every four days (on the same days as we conducted direct observations). The number of these brood items removed was the maximum that could be removed without disturbing the egg-cells, equating to 10-20 third/fourth-instar larvae or pupae per colony during each removal. We then divided the removed brood into two sets of equal size and placed each half into another randomly-selected experimental colony. This ensured that all R and C colonies contained third/fourth-instar larvae and pupae and that these brood items had a common origin (i.e. a different C or source colony).

### Collection of queens for mRNA-seq

To prepare for sampling a subset of queens for RNA extraction, at the start of the experiment we randomly assigned each queen to two sub-groups within each treatment, G1 (N = 20) and G2 (N = 15). (The additional R colony was not assigned to either sub-group or sampled for RNA.) Within each treatment, at TP1, when 10% (2/20) of the G1 queens had died, we randomly selected and removed six queens (termed TP1G queens) from the remaining G1 queens, flash-froze them in liquid nitrogen and stored them at −80°C. Similarly, at TP2, when 60% (9/15) of G2 queens had died, we randomly sampled the remaining six queens (termed TP2G queens) in G2 in the same way. The 10% and 60% mortality thresholds were selected to represent time-points marking the occurrence of low and high mortality, respectively, during the *B. terrestris* queen lifespan. We used relative time-points^61,62^, not ones based on absolute time, to account for potentially different ageing rates across the treatments (caused by differently shaped mortality curves if H1 is true) and to facilitate comparisons with other species. Hence, using this approach, we collected queens for RNA extraction at two time-points within each treatment (corresponding to the points of 10% (TP1) and 60% (TP2) queen mortality). The non-sampled G1 and G2 queens (termed life-history queens) were used to provide the queen longevity data, and all queens were used for egg-count and behavioural data until they died or were sampled (Table S28). Therefore, final queen sample sizes were: R:TP1G (N=6), R:TP2G (N=6), R:life-history (N=24); C:TP1G (N=6), C:TP2G (N=6), C:life-history (N=22) (Table S28).

### Tissue dissections, RNA extraction, and mRNA-seq

For each TP1G and TP2G queen, we dissected out the brain, fat body, and ovaries and flash froze all tissues in liquid nitrogen, storing each tissue from each queen individually at −80°C for later RNA extraction. We also measured the length of the marginal cell in each forewing (and calculated the mean marginal cell length) as an index of body size^63^ in all queens (except Q57) using a Zeiss Discovery v12 Stereo microscope (Zeiss, Oberkochen, Germany) with Axiovision software (Zeiss).

For RNA extraction we first fragmented the tissue in liquid nitrogen using a micropestle. We then added Tri-reagent (Sigma-Aldrich, Gillingham, Dorset, UK) at the level of 1 ml for each estimated 100 μg of tissue. We extracted RNA using the Direct-zol™ RNA extraction kit (Zymo Research, Irvine, CA, USA) according to the manufacturer’s protocol. We also performed an additional DNase treatment using the Turbo™ DNA-free kit (Thermo Fisher Scientific, Loughborough, UK) according to the manufacturer’s protocol. There was insufficient RNA for mRNA-seq from one TP2G brain sample in each treatment, so these two samples were excluded from further analysis. Using these procedures, we generated six biological replicates for each of the three tissues, two RNA sub-groups (TP1G and TP2G), and two treatments (R and C), excepting two TP2G brain samples for which there were 5 biological replicates, leading to a total of 72 (6 × 3 × 2 × 2) minus 2 = 70 RNA samples in total (Table S28). These were sent to a sequencing provider (Edinburgh Genomics) for Illumina 100 base pair, paired-end sequencing on two lanes of a NovaSeq6000 sequencer, creating two technical replicates for each biological replicate.

### Statistical analysis

All statistical analyses were conducted with the R (version 4.0.1) statistical programming platform in Rstudio^64^, using the ‘stats’, ‘lme4’, ‘glmmTMB’, and ‘survival’ packages. We used generalised linear mixed models (glmms) to investigate the effects of treatment on queen and colony fertility, longevity, worker egg-laying, worker aggression, and queen activity. We created versions of each model with individual fixed (and random effects where appropriate) included or excluded and used the Akaike Information Criterion (AIC) to determine the final model with the best fit to the data. To determine the significance of the fixed effects of all glmms, we compared the final model with a null model (all fixed effects removed) using a likelihood ratio test (Table S29). To conduct the glmms with negative binomial error distribution we used the ‘nb.glmer’ function in lme4, and to conduct glmms with Poisson error distribution for zero-inflated count data we used the glmmTMB function in the glmmTMB package. All other glmms were conducted with the lme4 package in R. We used a Cox’s proportional hazards survival analysis using the ‘coxph’ and ‘coxme’ functions in the ‘survival’ and ‘coxme’ packages to determine the effect of treatment on queen longevity for the life-history queens (i.e. censoring TP1G queens, TP2G queens, and Q57 in the models). Further details of the statistical analyses are reported in the Supplementary methods (*Statistical analysis of life-history data*).

### Bioinformatic analysis

We used several complementary approaches to assess the quality of the mRNA-seq reads and align them to the *B. terrestris* genome (Bombus_terrestris.Bter_1.0.dna.toplevel.fa^65^; Figures S15 - S20; Tables S2 - S11; see Supplementary methods for further details). A total of 68 samples met the quality threshold and were analysed, yielding final queen sample sizes for the mRNA-seq differential gene expression analysis as follows: brain: R:TP1G (N=6), R:TP2G (N=5); C:TP1G (N=6), C:TP2G (N=5); fat body: R:TP1G (N=6), R:TP2G (N=6); C:TP1G (N=6), C:TP2G (N=4); ovaries: R:TP1G (N=6), R:TP2G (N=6); C:TP1G (N=6), C:TP2G (N=6).

#### Differential gene expression between time-points within treatments

We summarised Kallisto pseudoalignments of mRNA-seq reads per gene (Table S3) to provide estimated counts for differential expression analysis in DESeq2 using an FDR adjusted *p*-value threshold of 0.05 (as detailed in *Quality assessment of mRNA-seq reads,* Supplementary methods). This procedure generated four lists of differentially expressed genes (DEGs) from the differential expression analysis for each tissue: genes more highly expressed in TP2 than TP1 (up-regulated genes) and genes more highly expressed in TP1 than TP2 (down-regulated genes) for both R and C treatments (Tables S4 – S6).

#### Differential gene expression comparison between time-points across treatments (i.e. comparison between relative and chronological age)

As R:TP2 and C:TP1 queens were sampled at similar chronological ages, we also compared gene expression profiles between R and C queens. We conducted differential gene expression analyses in DESeq2 using an FDR adjusted *p-*value threshold of 0.05 (as detailed in *Quality assessment of mRNA-seq reads,* Supplementary methods). For the analysis of R:TP2 and C:TP1, this procedure generated two lists of DEGs within each tissue: genes more highly expressed in C:TP1 than R:TP2 (up-regulated genes, i.e. increasing expression with chronological age) and genes more highly expressed in R:TP2 than C:TP1 (down-regulated genes, i.e. decreasing expression with chronological age). We then repeated this procedure to compare R:TP1 with C:TP1, and R:TP2 with C:TP2. Overall, these analyses resulted in 18 gene lists (3 comparisons × 3 tissues × 2 up-regulated/down-regulated gene lists) (Tables S23 - S25). We performed Fisher’s exact tests to detect significant overlaps between each pair of gene lists that overlapped in time-period. We adjusted for multiple testing using Bonferroni correction. To determine whether the seven genes that were differentially expressed between R:TP2 and C:TP1 were in the TI-J-LiFe network and enzymatic antioxidant gene set genes, we used OrthoFinder to isolate single-copy orthologues of these genes in *D. melanogaster*.

#### Queen age-related gene expression between treatments

We performed Fisher’s exact tests to detect significant overlaps between each pair of lists from R and C colonies for each tissue (brain, fat body and ovaries) and each direction of differential expression with respect to age (up- and down-regulated with age), resulting in six comparisons in total. The tests were performed using custom R (v4.0.1) scripts (R core team 2020) and adjusted for multiple testing using Bonferroni correction.

#### Comparative analysis of age-related gene expression with other insects

We compared lists of age-related genes in *B. terrestris* queens from the analysis of differential gene expression between time-points within treatments (this study) with lists of age- or ageing-related genes reported from other insects. For comparison with solitary insects, we used gene lists from two *D. melanogaster* mRNA-seq studies in which relative mortalities at sampling were comparable to those in the current study and two of the same tissues were studied, i.e. brain^42^ and fat body^41,42^. For comparisons with these studies, we downloaded the total gene list and the list of genes that were differentially expressed (as defined by each study) between the appropriate time-points. Pacifico et al.^42^ sampled at four time-points, so we used for the comparisons genes differentially expressed between 5 day old and 30 day old adults, as relative mortalities at these ages (5 days: ~30% mortality, 30 days: ~55% mortality) were very broadly similar to those in the current study. Chen et al.^41^ sampled at two time-points, so we used genes differentially expressed between these points (5 days post eclosion, 50 days post eclosion). We also compared the DEGs from the current study to the *D. melanogaster* genes in the GenAge database^43^, which catalogues experimentally validated, ageing-related genes. For comparisons of the current gene lists with those from other eusocial insects, available studies were as follows: a study showing that expression of *vitellogenin* increases with age in the head of *A. mellifera* queens^66^; a whole-body mRNA-seq study of the ant *C. obscurior* that isolated 12 genes showing an age-related expression pattern opposite to that of their *D. melanogaster* orthologues^28^, with eight of these 12 genes having single-copy orthologues with *B. terrestris;* and the two recent studies hypothesising that genes from the TI-J-LiFe network^33^ and enzymatic antioxidant gene set^35^ play a role in ageing in eusocial insects (see Introduction).

For all gene-list comparisons in these analyses, we identified single-copy orthologues of the relevant genes between *B. terrestris* and the comparison insect species using the OrthoFinder results (see *Gene Ontology (GO) enrichment analysis,* Supplementary methods). We then performed Fisher’s exact tests to detect significant overlaps between lists of DEGs from *B. terrestris* R or C queens (current study) and the lists of genes from the comparison species/studies. We compared the DEGs from Pacifico et al.^42^ and Chen et al.^41^ to the DEGs from *B. terrestris* R and C queens in brain and fat body (respectively), comparing between lists of genes differentially expressed in the same direction with respect to time (2 treatments × 2 tissues × 2 directions of differential expression = total of 8 comparisons). We compared the GenAge gene list to the combined up- and down-regulated genes within each treatment and tissue in the current study, as there was no expectation as to whether the GenAge genes would be up- or down-regulated with age (total 6 comparisons). The small number of genes available for comparison from *C. obscurior* (8) led us to compare whether *B. terrestris* R and C DEGs matched *C. obscurior* or *D. melanogaster* expression patterns of these genes, rather than whether the list significantly overlapped with the DEGs from R and C (total 6 comparisons). We modified the approach taken by Korb et al.^33^ to compare the DEGs from *B. terrestris* (current study) to the lists of *D. melanogaster* genes associated with the TI-J-LiFe network and the enzymatic antioxidant gene set. For this, we combined up- and down-regulated DEGs within each treatment and tissue in the current study and ranked them by log fold-change in expression with time. We then picked out the 50 genes with the most positive log-fold change and the 50 genes with the most negative log fold-change within each treatment/tissue (hereafter, ‘top ± 50 genes’). Likewise, where the number of DEGs in a treatment/tissue allowed, we extracted the top ± 100 genes, top ±200 genes, top ± 300 genes, top ± 500 genes, and all genes. We then compared the gene lists from the TI-J-LiFe network and the enzymatic antioxidant gene set to the available ‘top ±’ genes lists for each treatment/tissue combination (total of 24 comparisons for each of the TI-J-LiFe network and the enzymatic antioxidant gene set gene lists). As before, we performed Fisher’s exact tests using custom R (v4.0.1) scripts^64^ and adjusted for multiple testing using Bonferroni correction.

## Data availability

Data on life history, behaviour, morphometrics, etc., from this study are available in Collins et al.^67^. The raw mRNA-seq sequencing data have been deposited in the National Center for Biotechnology Information’s (NCBI’s) Gene Expression Omnibus (GEO) at https://www.ncbi.nlm.nih.gov/geo/ and are available under series accession number *GSE172422.* The code used for the analyses is available at https://github.com/davidprince84/Collins-et-al_NE-R000875-1_obj1_mRNA-seq_scripts.

## Acknowledgements

We thank Ryan Brock, Liam Crowther, and Jenny Livesey for helpful comments on the experimental design and statistical analysis and/or for assistance with maintaining bee colonies. We also thank Annette Van Oystaeyen (Biobest Group NV) for comments on the manuscript and for invaluable help in supplying young *B. terrestris* colonies. This work was funded by the UK’s Natural Environment Research Council (NERC research grant reference number NE/R000875/1). Laboratory work (library preparation and sequencing) was supported and performed by the NERC Biomolecular Analysis Facility (NBAF) at the University of Edinburgh (Edinburgh Genomics). Edinburgh Genomics is partly supported through core grants from BBSRC (BB/T017864/1) and NERC (UKSBS PR18037). The gene expression analysis was carried out on the High Performance Computing Cluster supported by the Research and Specialist Computing Support service at the University of East Anglia.

## Author Contributions

D.H.C., D.C.P., T.C., and A.F.G.B. designed the study. D.H.C. and J.L.D. performed the experimental manipulations, extracted the RNA samples, and recorded the life-history data. D.H.C. analysed the life-history data and D.C.P. analysed the mRNA-seq data. D.H.C., D.C.P., and A.F.G.B. wrote the manuscript and all authors commented on drafts. T.C. and A.F.G.B raised funding.

