## Supplementary text and figures for "Eusocial insect queens show costs of reproduction and transcriptomic signatures of reduced longevity"

### Supplementary information

#### Supplementary Methods

##### *Statistical analyses of life-history data*

We used two indices of queen fertility (baseline queen fertility and treatment queen fertility, see *Experimental manipulation and colony fertility*, Methods, main text) to test for the effect of treatment on queen fertility. The model consisted of a glmm with negative binomial error distribution to account for overdispersion. Treatment and a binary variable (representing the periods before and after experimental manipulations were started) were fitted as fixed effects and ID (the unique number given to each queen/colony; Table S28) was fitted as a random effect. We also used a glmm with negative binomial error distribution to test the effect of treatment on colony fertility (i.e. the unique egg count for each colony, including worker-laid eggs, on each count day until the last queen death on day 158) as a function of time, with treatment and experimental day (fitted as a quadratic term to account for its non-linear effect on fertility) as fixed effects in the model. We created versions of the model that included random effects for ID and experimental day. Each model was compared with and without these terms using the Akaike Information Criterion (AIC) to determine the best model fit. The model with the lowest AIC included ID and experimental day as random effects (Table S29).

To test whether colony fertility was affected by worker egg-laying, we used zero-inflated glmms with binomial error distribution to determine whether there were any differences in the levels of observed worker egg-laying between R and C colonies. The data were not found to be overdispersed. We fitted treatment and day as fixed effects and we created versions of the model that included random effects for ID and experimental day. The model was compared with and without these terms using AIC to determine the best model fit. The model with the lowest AIC included ID and day as random effects (Table S29). We did not statistically analyse the number of filmed worker egg-laying events, as only nine workers were observed laying eggs in the total 32 hours of analysed digital film.

We used a negative binomial glmm to determine whether the following variables (all expressed in units of numbers of workers) were significantly different between treatments to control for their potential influence on queen longevity (as defined in *Queen longevity*, Methods, main text): callow workers added, excess workers removed, dead workers removed (each as defined in *Colony maintenance*, Methods, main text), egg-laying workers removed, and aggressive workers removed (each as defined in *Observed worker aggression and observed worker egg-laying*, Methods, main text). For each of these variables, including age (the number of days between day 1 and queen death) as a fixed effect significantly improved the model fit (lower AIC, likelihood ratio test $p < 0.05$), so it was included in the final model (Table S29).

We used a Cox's proportional hazards survival analysis to determine the effect of treatment on queen longevity. Life-history queens were included in the model, and the queens sampled for RNA (TP1G and TP2G queens) and Q57 were included as censored individuals. We used graphical and analytical tests to ensure there were no violations of the proportional hazards assumption. As queen fertility (Figures 2a, 2b), observed worker egg-laying (Figure 2c), excess workers removed (see *Colony maintenance*, Methods, main text) and egg-laying workers removed (see *Observed worker aggression and observed worker egg laying*, Methods, main text) were significantly different between R and C colonies, we included these effects as additional fixed effects to control for their effect on queen longevity. Although observed worker aggression was not significantly different between R and C colonies (Figure 3c), we also included this as an additional fixed effect. To account for larger numbers of observations for older colonies, each variable was converted into a per-colony rate (except for queen fertility, which was already calculated as a per-colony rate – see *Experimental manipulation and colony fertility*, Methods, main text). The per-colony rates were calculated by summing all values for the given variable across the total number of observation periods and then dividing by the total number of observation periods for each colony. We then produced one iteration for each variant of the Cox's proportional hazards model of queen longevity where each per-colony-rate (queen fertility, observed worker egg-laying, observed worker aggression, excess workers removed, and egg-laying workers removed) was included as an additional fixed effect to control for its effects on queen longevity. We also compared the model with a model in which marginal cell width was included as a fixed effect to control for the potential effect of body size on queen longevity. The model was compared with and without all of these terms using AIC to determine the best model fit. Independently adding excess workers removed as a fixed effect, but not the other variables (marginal cell width, queen fertility, observed worker egg-laying, observed worker aggression, egg-laying and workers removed), significantly improved the model fit (lower AIC value, likelihood ratio test $p < 0.05$) compared to the model with all these variables removed. This meant that, independently of treatment, excess workers removed was a significant predictor of queen longevity, with a higher rate of excess workers removed being associated with reduced queen longevity (Cox's proportional hazards analysis: hazards ratio = 1.797, $z = 4.903$, $p < 0.001$). Therefore, the final reported Cox's proportional hazards model of queen longevity included treatment and excess workers removed as fixed effects and none of the other variables were included in the final model (Table S29).

To further test whether queen longevity was affected by worker aggression, we used zero-inflated glmms with Poisson error distribution to determine whether there were any differences in the levels of observed worker aggression between R and C colonies. As with observed worker egg-laying, we fitted treatment and day as fixed effects and we created versions of the model that included random effects for ID, experimental day, and treatment. The model was compared with and without these terms using AIC to determine the best model fit. The model with the lowest AIC included ID and day as random effects. We also tested whether there were any differences in the levels of filmed worker aggression between the R and C colonies for which we had film data. As filmed aggression rates were low (0.5 incidences per colony per hour of digital film), we analysed filmed worker aggression as a binary index indicating whether aggression was observed/not observed during each film period in each colony. We then used a glmm with binomial error distribution to test whether treatment and film period affected the filmed worker aggression index within colonies. In this model, colony ID was fitted as a random effect (Table S29).

We used glmms with binomial error distributions to determine whether there were any differences in observed queen activity, which was fitted as a binary response variable. We fitted treatment and day as fixed effects and we created versions of the model that included random effects for ID, experimental day, and treatment. Each model was compared with and without these terms using AIC to determine the best model fit. The model with the lowest AIC included ID and day as random effects in both cases. To test whether there were any differences in filmed queen activity, we used a glmm with binomial error distribution with the number of seconds a queen spent active, relative to total number of seconds she was filmed during each film period, fitted as the response variable, and treatment and film period fitted as fixed effects. We created versions of each model that included random effects for ID, film period, and treatment. The model with the lowest AIC included ID as a random effect (Table S29).

To test whether there was significant relationship between queen fertility and queen longevity within each treatment (Figure S1), we used an ANCOVA to determine the combined effects of treatment and longevity on queen fertility after treatment (between days 5-25 inclusive). We used graphical and analytical tests to ensure that there were no violations of the ANCOVA assumptions of homogeneity of variance (Levene's test, $p = 0.115$) and normality of residuals (Shapiro-Wilk test, $p = 0.410$).

*Quality assessment of mRNA-seq reads*

We used several complementary approaches to assess the quality of the mRNA-seq reads. First, we used FastQC v0.11.9[1] to examine a range of quality measures including base quality and potential adapter contamination in each sample, with the results for each sample combined into a report for each tissue (brain, fat body, and ovaries) using the MultiQC v1.9 python library[2] with Python v3.7[3] (Supplementary files S1 - S3). Subsequently, we aligned reads against the *Bombus terrestris* genome (Bombus_terrestris.Bter_1.0.dna.toplevel.fa) using HISAT2 v2.1.0[4] and recorded mapping statistics (Table S8). We used the HISAT2 alignment files to assess gene body coverage and junction saturation using the RSeQC v3.0.1 Python library[5] with Python v3.7 and to assess which genomic features the reads aligned to using the ALFA v1.1.1 python library[6] with Python v3.7. These approaches showed that some samples in each tissue had large numbers of overrepresented sequences and atypical per-sequence GC content (as assessed with FastQC), and low read alignment (6.4 – 69.5%) (as assessed with HISAT2). We examined several of the overrepresented sequences with BLAST (https://blast.ncbi.nlm.nih.gov/Blast.cgi)[7], using blastn against the nr/nt database, which suggested that the sequences originated from bee viruses.

To further investigate the identity of reads not mapping to the *B. terrestris* genome, we pseudoaligned reads to the Holobee database (HB_Bar_v2016.1) (https://data.nal.usda.gov/dataset/holobee-database-v20161) with Kallisto v0.46.1[8]. This is a curated database of publicly accessioned nucleotide sequences from honeybee (*Apis mellifera*) holobionts (no similar resource exists for bumblebees, which belong to the same family, the Apidae, as *Apis*). Examining the normalised read counts revealed 10 sequences, from 8 different holobionts, that had >500 total counts in at least 2 out of the 3 tissues (Tables S9 – S11). Plotting the normalised counts against the percentage of reads aligned to the *B. terrestris* genome with HISAT2 for each sample in fat body revealed a negative relationship in two holobiont sequences, both from slow bee paralysis virus (SBPV) (Figure S15). These results imply that, in these libraries, large numbers of SBPV reads were present that caused low percentages of reads to align to the *B. terrestris* genome. The same trend was seen in brain and to a lesser extent in ovaries (Figure S16). Overall, SBPV viral RNA was observed in 11/23 brain samples, 12/24 fat body samples, and 12/24 ovaries samples. In each case, if a given queen showed SBPV viral RNA sequences to be present in one tissue, these sequences were also present in the other two tissues sampled, suggesting that 12 queens in total were infected with SBPV (with one such queen not yielding a brain sample that underwent mRNA-seq). Of the 12 infected queens, 4 were Removal (R) queens and 8 were Control (C) queens. We assessed whether the presence of SBPV was likely to significantly alter gene expression in the samples by examining sample clustering, based on gene expression, using a principal component analysis (PCA). We pseudoaligned reads to the *B. terrestris* transcriptome (Bombus_terrestris.Bter_1.0.cdna.all.fa) with Kallisto v0.46.1[8], with estimated transcript counts being summarised per gene with tximport v1.16.1[9], followed by differential expression analysis and data visualisation using DESeq2 v1.28.1[10]. First, we assessed the variation in gene expression between technical replicates of the same biological replicate and found that technical replicates were extremely similar (data not shown). Therefore, in all subsequent analyses, we collapsed technical replicates using the collapseReplicates() function in DESeq2. The subsequent PCA plot revealed that samples in each tissue clustered by treatment and/or time-point, rather than by presence of SBPV (Figure S17), suggesting that the presence of SBPV was unlikely to significantly alter gene expression in the samples. Accordingly, we did not exclude any samples from the analysis on the basis of SBPV presence. However, we included in the gene expression analyses only samples exceeding a minimum threshold of 12 million pseudoaligned read pairs per sample (from the two technical replicates combined), so prioritising biological replication over sequencing depth as being more informative in such analyses[11,12]. Two fat body samples (C TP2G biological replicates 4 and 6) did not meet this threshold and so we excluded them from the subsequent analysis.

*Differential expression analysis*

We performed differential expression analysis using DESeq2 with an FDR adjusted *p*-value threshold of 0.05 and the model ~ virus + condition, in which virus was a categorical factor denoting the presence of many SBPV-aligning reads in a sample and condition was a categorical factor denoting the combined treatment and time point of a sample. We produced boxplots of the normalised count data and principal component analysis from DESeq2 for each tissue to check normalisation and library clustering, respectively (Figures S18 – S20).

*Gene Ontology (GO) enrichment analysis*

To perform GO enrichment analysis, and comparative analyses with other insect species, we used Orthofinder V2.5.2[13] to identify orthologues between *B. terrestris*, *Drosophila melanogaster*, *A. mellifera*, and *Cardiocondyla obscurior*. *D. melanogaster* single-copy orthologues for *B. terrestris* differentially expressed genes (DEGs) were used for GO enrichment analysis, as GO annotations for *D. melanogaster* are much more complete. GO enrichment analysis was then performed in R (v4.0.1)[14] via the clusterProfiler package (v3.16.1)[15] using biological processes GO annotations from the org.Dm.eg.db package (v3.11.4)[16]. We used an over-representation test[17] to identify GO terms that were significantly overrepresented ($p < 0.05$ after adjustment for multiple testing with *Benjamini-Hochberg*) in a set of DEGs against a background consisting of all genes that were expressed in the relevant tissue. Redundancy in the resulting enriched GO terms was reduced using the GoSemSim package (v2.14.2)[18,19].

#### Supplementary Results

*Queen fertility, colony fertility and worker egg-laying*

From day 26 (first recorded worker egg-laying in any colony), some differences in treatment-specific colony fertility were possibly due to worker egg-laying. Consistent with this, after day 26, R colonies had significantly higher levels of observed worker egg-laying than C colonies (zero-inflated negative binomial glmm: Treatment: b = 2.917, SEb = 0.831, z =

3.509, $p < 0.001$; Figure 2c). Observed worker egg-laying was also significantly time-dependent, peaking at day 76 and then declining for the rest of the experiment (Time: $b = 40.156$, $SEb = 9.133$, $z = 4.397$, $p < 0.001$; Figure 2c). As R queens showed significantly higher fertility than C queens before day 26 (see Results, main text, for details), and as colony fertility peaked on day 35 (Figure 2b) whereas observed worker egg-laying did not peak until day 76, these results show that, as intended, the R treatment increased queen egg-laying rate and that this occurred both before and after the start of worker egg-laying.

*Queen activity and response to disturbance*

There was no significant difference in observed queen activity (binomial glmm: $b = -0.176$, $SEb = 0.419$, $z = -0.419$, $p = 0.675$; Figure S6), filmed queen activity (binomial glmm: $b = -0.067$, $SEb = 0.297$, $z = 0.224$, $p = 0.822$; Figure S7), or queens' response to disturbance ($b = -3.3$, $SEb = 1.97$, $z = -1.678$, $p = 0.093$; Figure S8) between R and C queens. Hence the decreased longevity observed in R queens was not caused by differences in queen activity levels or responsiveness between treatments.

*Read counts and DEGs in mRNA-seq libraries*

Across the 70 libraries, mRNA-seq resulted in a mean of 55,280,953 read pairs per library for brain, 61,864,754 read pairs per library for fat body and 69,025,352 reads pairs per library for ovaries (Table S2). The libraries pseudoaligned to the *B. terrestris* transcriptome with a mean percentage pseudoalignment of 77.3% (range 49.4 – 88.4%) for brain, 58.7% (5.5 – 90.0%) for fat body (including the two fat body libraries that were excluded from further analysis), and 75.9% (32.7 – 83.1%) for ovaries (Table S3).

In total, between the two time-points (TP1 and TP2), and pooling across both treatments (R and C), there were 836 DEGs in brain, 2,572 DEGs in fat body and 6,440 DEGs in ovaries. In brain, in R queens: 30 genes were more expressed in TP2G than TP1G (i.e. genes which increased in expression with age; herein defined as 'up-regulated genes') and 7 genes were more expressed in TP1G than TP2G (i.e. genes which decreased in expression with age; herein defined as 'down-regulated genes'); in C queens: 482 genes were up-regulated and 317 genes were down-regulated (Figures 6a, S9; Table S4). In fat body, in R queens: 430 genes were up-regulated and 412 genes were down-regulated; in C queens: 923 genes were up-regulated and 807 genes were down-regulated (Figures 6b, S10; Table S5). In ovaries, in R queens: 3 genes were up-regulated and 0 genes were down-regulated; in C queens: 3,520 genes were up-regulated and 2,917 genes were down-regulated (Figures 6c, S11; Table S6).

*Gene Ontology (GO) enrichment analysis*

OrthoFinder identified 6,074 single-copy orthologues between *B. terrestris* and *D. melanogaster* (57.4% of the 10,591 genes expressed in the *B. terrestris* mRNA-seq libraries). These were used to isolate 248 nonredundant enriched gene ontology (GO) terms for the DEGs (Table S7).

In brain, up-regulated DEGs in R queens were not enriched for GO terms, while up-regulated DEGs in C queens were enriched for GO terms in cytoplasmic translation (GO:0002181). Down-regulated DEGs in R queens were enriched for GO terms associated with ion transport (4/9 terms) and amino acid processes (3/9 terms), while down-regulated DEGs in C queens were not enriched for GO terms (Table S7).

In fat body, up-regulated DEGs in R queens were enriched for GO terms associated with DNA recombination (GO:0006310) and cellular response to DNA damage stimulus (GO:0006974), while up-regulated DEGs in C queens were enriched for a variety of processes with no obvious pattern. Down-regulated DEGs in R queens were enriched for GO terms associated with endoplasmic reticulum, Golgi, and vesicle trafficking (7/14 terms), while down-regulated DEGs in C queens were enriched for GO terms associated with metabolic and biosynthetic processes (7/10 terms) (Table S7).

In ovaries, in R queens, GO analysis was not conducted as up- and down-regulated genes comprised 3 and 0 DEGs, respectively. Up-regulated DEGs in C queens were enriched for GO terms associated with development (19/97 terms referred to 'development'). Down-regulated DEGs in C queens were enriched for GO terms associated with regulation of the cell cycle and associated nuclear and chromosomal changes (18/45 terms) (Table S7).

*DEGs between overlapping time periods*

There were significant and high percentage overlaps in DEGs between time-periods (intervals between sampling time-points) that overlapped in chronological time.

The overlaps of DEGs between R:TP1 and R:TP2 versus between R:TP1 and C:TP1 were: up-regulated: brain: 26/30 (86.3%) DEGs from R:TP1 to R:TP2 overlapped with 143 DEGs from R:TP1 to C:TP1 (Fisher's exact test: $p < 0.001$); fat body: 385/430 (89.5%) DEGs from

R:TP1 to R:TP2 overlapped with 1077 DEGs from R:TP1 to C:TP1 (Fisher's exact test: $p < 0.001$); ovaries: 2/3 DEGs from R:TP1 to R:TP2 overlapped with 15 DEGs from R:TP1 to C:TP1, but gene numbers were too low for statistical comparison; down-regulated: brain: 3/7 DEGs from R:TP1 to R:TP2 overlapped with 96 DEGs from R:TP1 to C:TP1, but gene numbers were too low for statistical comparison; fat body: 353/412 (85.7%) DEGs from R:TP1 to R:TP2 overlapped with 986 DEGs from R:TP1 to C:TP1 (Fisher's exact test: $p < 0.001$); ovaries: no comparison was possible as there were 0 DEGs from R:TP1 to R:TP2) (Tables S26, S27).

The overlaps of DEGs between C:TP1 and C:TP2 versus between R:TP2 to C:TP2 were: up-regulated: brain: 362/482 (75.1%) DEGs from C:TP1 to C:TP2 overlapped with 728 DEGs from R:TP2 to C:TP2 (Fisher's exact test: $p < 0.001$); fat body: 757/923 (82%) DEGs from C:TP1 to C:TP2 overlapped with 1338 DEGs from R:TP2 to C:TP2 (Fisher's exact test: $p < 0.001$); ovaries: 3189/3520 (90.6%) DEGs from C:TP1 to C:TP2 overlapped with 3433 DEGs from R:TP2 to C:TP2 (Fisher's exact test: $p < 0.001$); down-regulated: brain: 239/317 (75.4%) DEGs from C:TP1 to C:TP2 overlapped with 645 DEGs from R:TP2 to C:TP2 (Fisher's exact test: $p < 0.001$); fat body: 614/807 (76.1%) DEGs from C:TP1 to C:TP2 overlapped with 1217 DEGs from R:TP2 to C:TP2 (Fisher's exact test: $p < 0.001$); ovaries: 2431/2917 (83.3%) DEGs from C:TP1 to C:TP2 overlapped with 2611 DEGs from R:TP2 to C:TP2 (Tables S26, S27).

#### Supplementary Discussion

##### *Worker-to-queen aggression*

Worker aggression directed at queens did not differ between treatments, and therefore the decreased longevity of R queens was not caused by worker aggression (Figures 3c, S4). Almond et al.[20] showed that egg removal caused an increase in worker-to-queen aggression in *B. terrestris*, which was interpreted as workers responding in a self-interested manner to a perceived loss of queen fecundity. However, the current experimental design differs from that of Almond et al.[20] in two key respects. First, colony size was maintained throughout at a constant level (20 workers), which may have affected workers' response to a perceived loss of queen fecundity. Second, we removed all aggressive workers from colonies in both treatments as and when they were detected (and replaced them with callow workers) (see

Methods, main text). This meant that individual workers in the current experiment would have been unable to maintain a sustained response to queen egg removal. Furthermore, Almond et al.[20] found that egg removal increased worker aggression in the pre-competition period only, whereas the current experiment extended far into the post-competition point period (as queens had a mean longevity of 89.4 days but the first competition point occurred on day 26).

R colonies had significantly more egg-laying workers removed than C colonies, and significantly fewer excess workers removed, which may have affected the within-colony worker age-structure within each treatment and so have influenced the results. However, colony age-structure and/or worker removal is unlikely to have accounted for the difference in longevity between R and C queens in the current experiment for the following reasons. First, the number of egg-laying workers removed was low (means of 2.3 and 1.5 workers removed, respectively, from R and C colonies) and therefore unlikely to have substantially altered worker age-structure given that each colony was maintained at a size of 20 workers. Accordingly, the number of egg-laying workers removed was not a significant predictor of queen longevity. Second, although there were significantly fewer excess workers removed from R colonies than C colonies (means of 39.1 and 67.9 workers removed, respectively), all the removed workers were callow workers, and their removal was therefore unlikely to have affected worker age-structure in either treatment. (Similarly, all added workers were callows, which again would not have altered the worker age-structure differentially.) Moreover, although excess workers removed was a significantly predictor of queen longevity (see *Statistical analysis of life-history females*, Supplementary Methods), it affected it negatively; hence, as R colonies had fewer excess workers removed, the effect of excess workers removed as a factor was counter to the prediction of H1, and could not have accounted for the independent effect of treatment in reducing overall queen longevity in R colonies.

*Gene expression differences between R and C queens*

In the current study, R and C queens differed in both their fertility and longevity, which suggests that some of the gene expression differences between the treatments (with relative age) could have been caused by fertility differences as well as R queens' increased costs of reproduction[21–24]. However, the greater fertility of R queens is unlikely to have accounted entirely for the treatment-specific differences in age-related gene expression. First, R queens overall showed decreased age-related gene expression differences compared to C queens

(Figure 4), which is unexplained if gene expression changes were primarily driven by the increased reproduction of R queens. Second, if gene expression changes were primarily driven by R queens' increased reproduction, we would have expected to see such changes mainly in fat body[24,25] and ovaries, but in fact such changes occurred mainly in brain and fat body, with very few changes occurring in ovaries (Figure 4).

#### Supplementary Figures

**Figure S1**

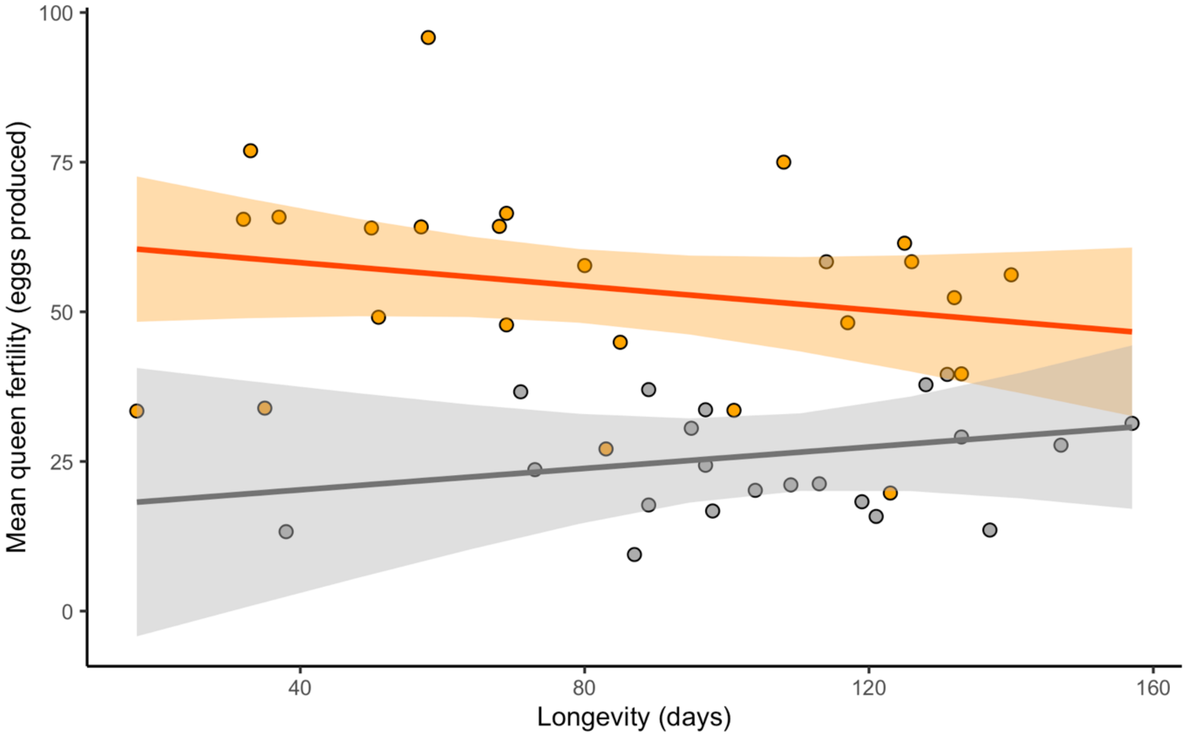

Figure S1. The relationship between queen longevity (days) and queen fertility during treatment (mean number of eggs laid per 48-hour period between days 5-25 inclusive) in R (eggs removed; orange circles, N = 24) and C (eggs removed and replaced; gray circles, N = 22) queens of *Bombus terrestris*. Regression line (and confidence intervals): the model fit from the ANCOVA analysis of these data (see *Statistical analyses of life-history data*, Supplementary methods). There was no significant relationship between fertility and longevity in either treatment (ANCOVA: R: $F = 0.669$, $df = 1, 21$, $p = 0.423$; C: $F = 0.868$, $df = 1, 19$, $p = 0.363$) and no evidence of an interaction of treatment and fertility in their effects on longevity across all queens (i.e. the slopes for R and C queens were not significantly different; ANCOVA: $F = 0.367$, $df = 1, 40$, $p = 0.548$). However, consistent with H1, there were non-significant trends for the longest-lived R queens to have the lowest average fertilities (negative fertility-longevity relationship) and the longest-lived C queens to have the highest average fertilities (positive fertility-longevity relationship).

**Figure S2**

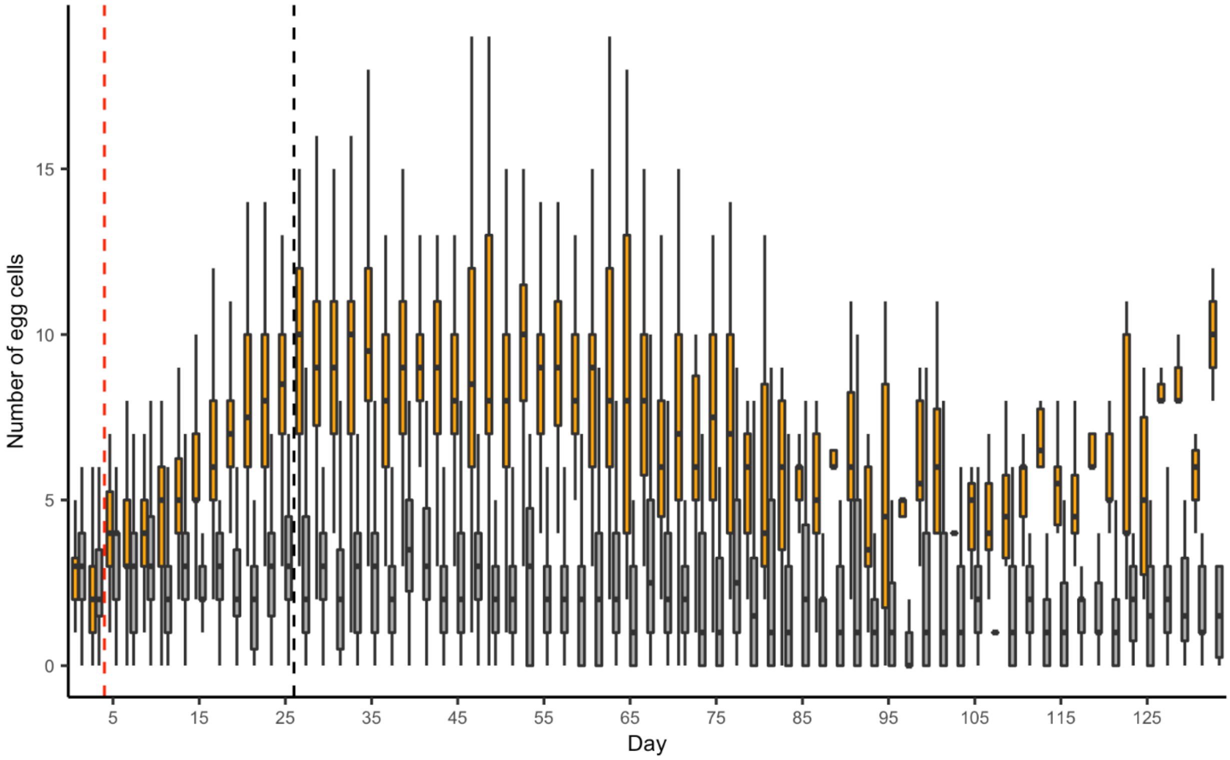

Figure S2. Total number of egg-cells for R (eggs removed; orange boxes; N = 36 on day 1 declining to N = 1 on day 134) and C (eggs removed and replaced; gray boxes; N = 35 on day 1 declining to N = 1 on day 148) *Bombus terrestris* queens/colonies until day 134. Vertical red dotted line: day when manipulations started; vertical black dotted line: day when worker egg-laying was first observed in any colony. Sample sizes on each day are in Table S28. Outliers are not shown. Black horizontal bars: median values for each treatment; boxes: interquartile ranges; whiskers: ranges up to 1.5 × the interquartile range. Over the course of the experiment, R colonies had significantly higher egg-cell numbers than C colonies (b = 1.107, SEb = 0.110, z = 19.690, $p < 0.001$).

**Figure S3**

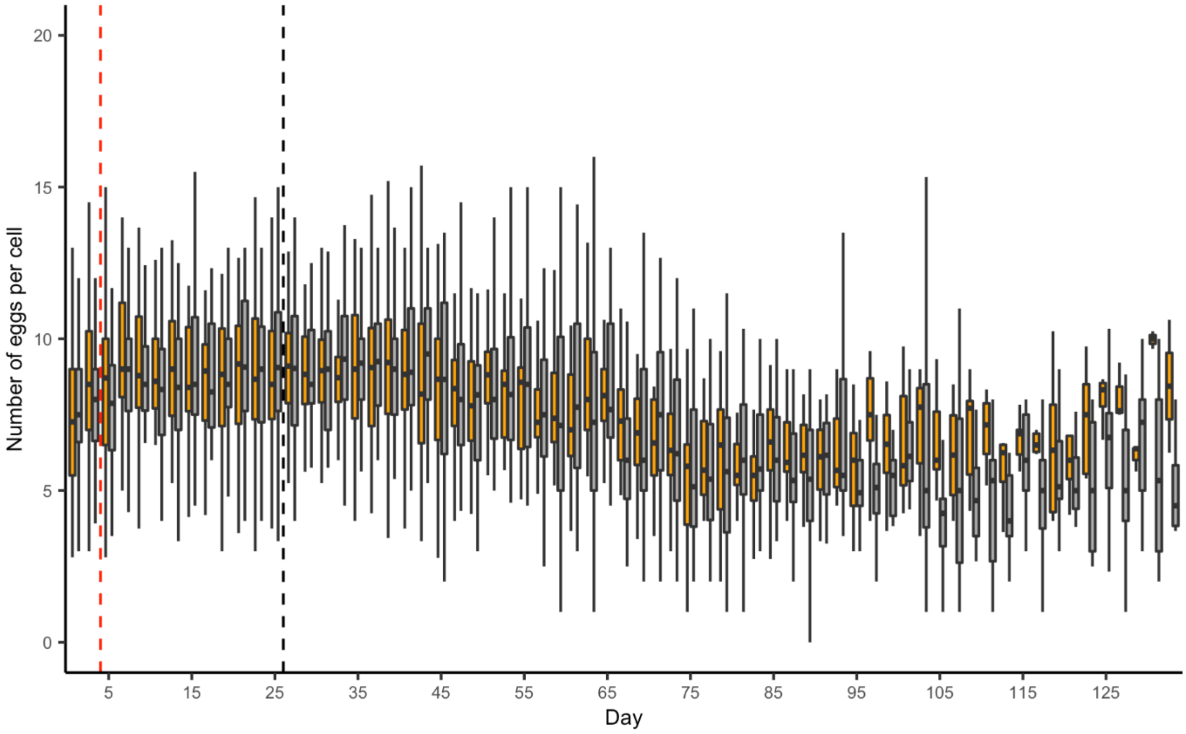

Figure S3. Mean number of eggs per egg-cell for R (eggs removed; orange boxes; N = 36 on day 1 declining to N = 1 on day 134) and C (eggs removed and replaced; gray boxes; N = 35 on day 1 declining to N = 1 on day 148) *Bombus terrestris* queens/colonies until day 134. Vertical red dotted line: day when manipulations started; vertical black dotted line: day when worker egg-laying was first observed in any colony. Sample sizes on each day are in Table S28. Outliers are not shown. Black horizontal bars: median values for each treatment; boxes: interquartile ranges; whiskers: ranges up to 1.5 × the interquartile range.

**Figure S4**

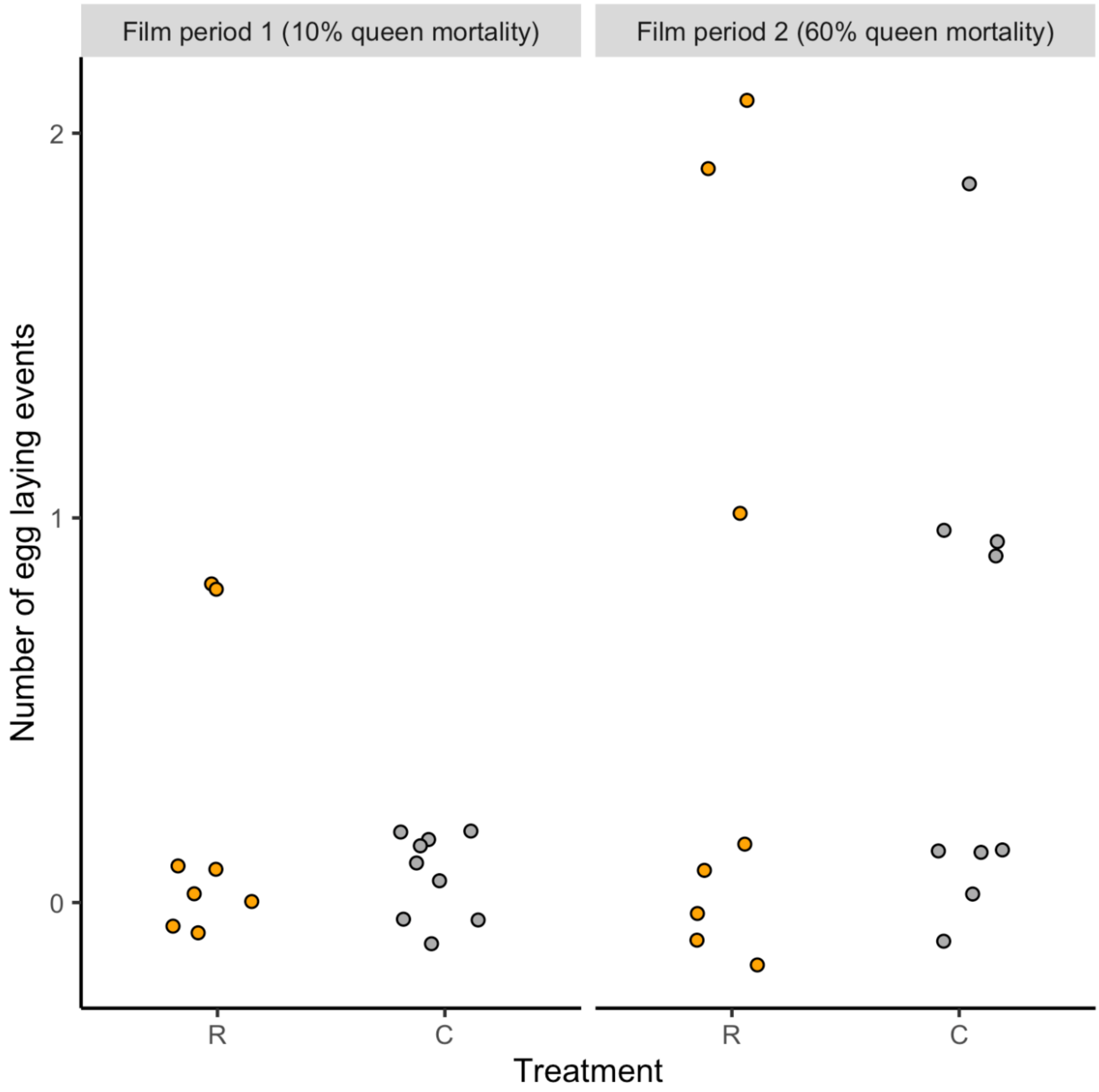

Figure S4. Filmed worker egg-laying in R (eggs removed; orange circles, N=8) and C (eggs removed and replaced; gray circles, N=9) colonies of *Bombus terrestris*. Filming was conducted during 2 × 1-hour film periods, i.e. film period 1 on day 48 and film period 2 on day 99, to coincide with time-point 1 (TP1, 10% queen mortality) and time-point 2 (TP2, 60% queen mortality) of the C colonies, respectively. Points are offset around each integral value on both axes. R colonies had higher rates of filmed worker egg-laying than C colonies; however, due to the low sample size for filmed worker egg-laying events this could not be analysed statistically.

**Figure S5**

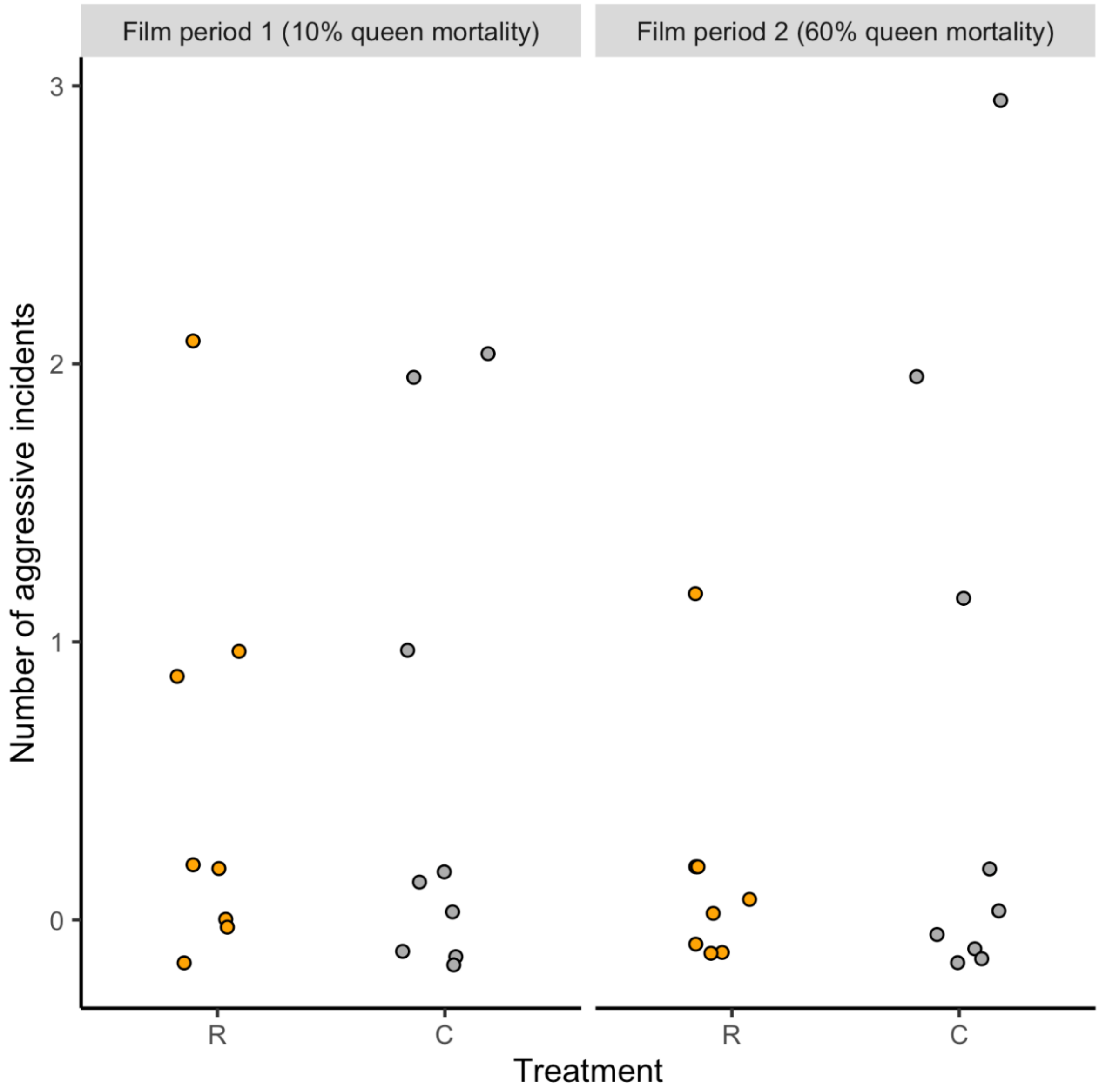

Figure S5. Number of worker-to-queen aggressive incidents in R (eggs removed; orange circles, N=8) and C (eggs removed and replaced; gray circles, N=9) colonies of *Bombus terrestris* during 2 × 1 hour-long film periods. Film periods were as defined in the Figure S4 legend. Points are offset around each integral value on both axes. There was no significant difference in filmed worker aggression between R and C colonies (see Results, main text, for details).

**Figure S6**

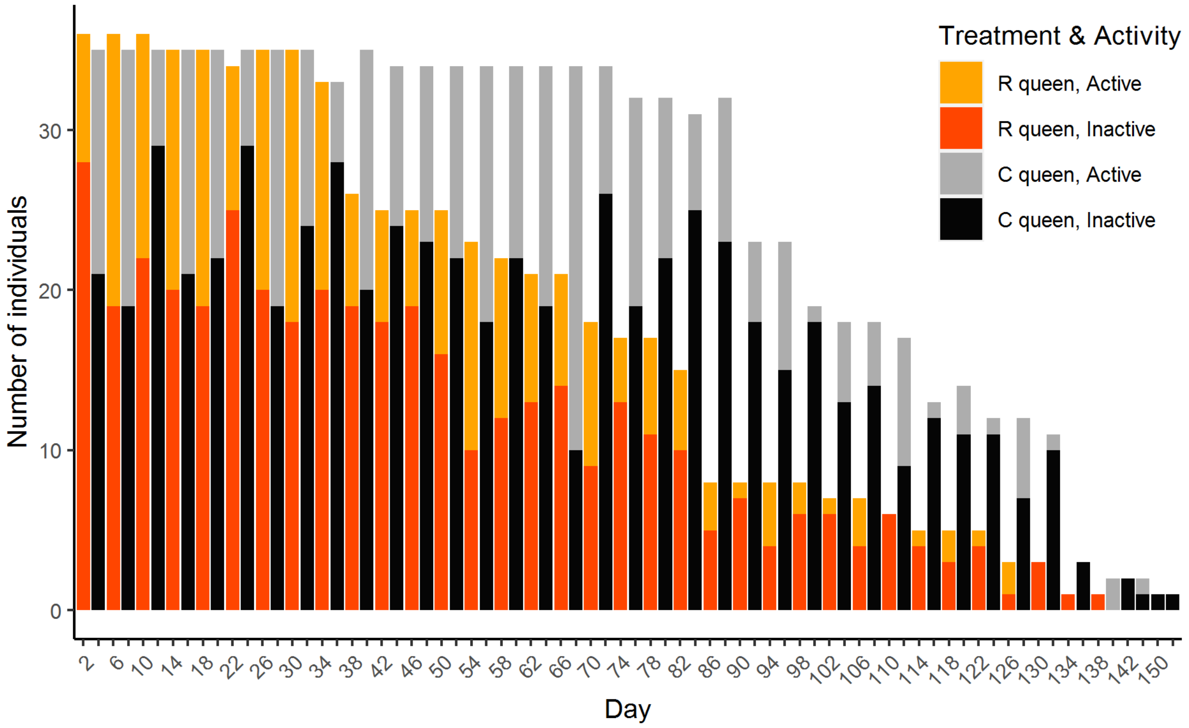

Figure S6. Observed activity levels for R (eggs removed; pale/dark orange bars, N = 36 on day 1 declining to N = 1 on day 134) and C (eggs removed and replaced; pale/dark gray bars, N = 36 on day 1 declining to N = 1 on day 148) queens of *Bombus terrestris* over the course of the experiment. Bar height: number of queens alive on each day in each treatment; pale relative to dark colour (within each bar): proportion of queens that were active (moving when recorded) to inactive (not moving when recorded) on each day in each treatment. There was no significant difference in observed activity levels between R and C colonies (see Supplementary Results ). In addition, the proportion of time spent active did not change throughout the experiment in either treatment (binomial glmm: b = -0.056, SEb = 0.110, z = -0.510, p = 0.610).

**Figure S7**

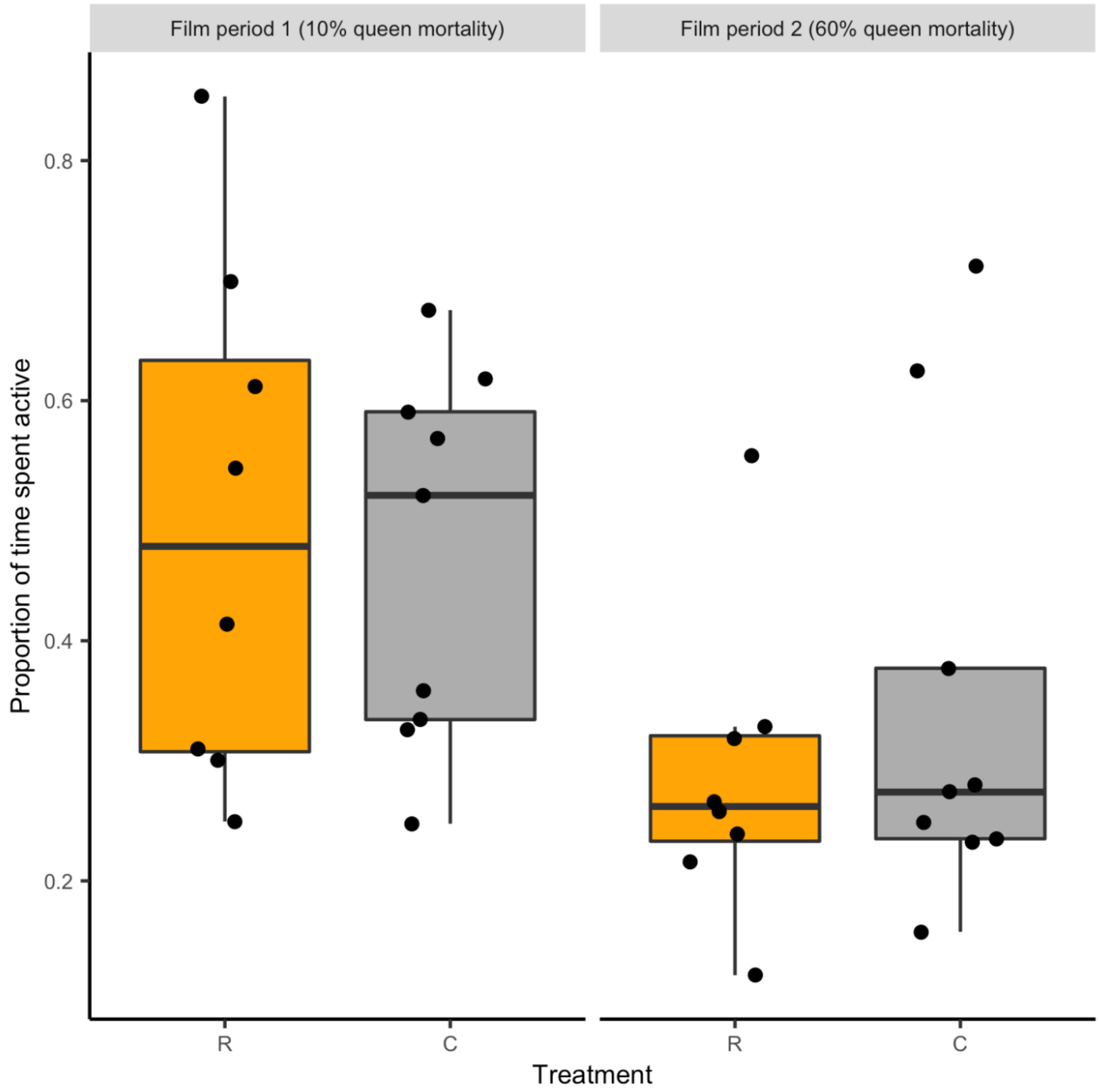

Figure S7. Proportion of time spent active for R (eggs removed; orange boxes, N = 8) and C (eggs removed and replaced; gray boxes, N = 9) *Bombus terrestris* queens during 2 × 1-hour film periods. Film periods were as defined in the Figure S4 legend. Black circles: individual data values; black horizontal bars: median values for each treatment, boxes: interquartile ranges; whiskers: ranges up to 1.5 × the interquartile range. The amount of filmed activity was significantly lower in the second than in the first period in both treatments (binomial glmm: $b = -0.922$, $SEb = 0.019$, $z = -48.830$, $p < 0.001$). However, there was no significant difference in filmed queen activity levels between R and C colonies independently of period (see Supplementary Results).

**Figure S8**

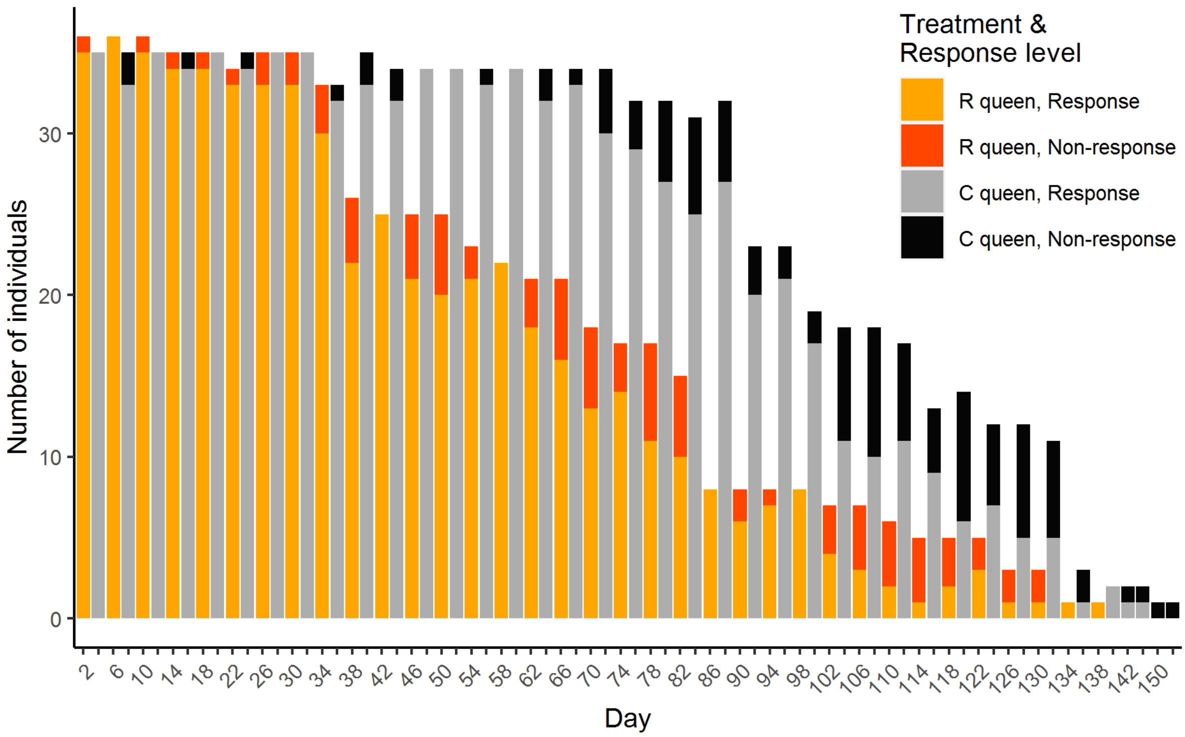

Figure S8. Observed response to disturbance for R (eggs removed; pale/dark orange bars, N = 36 on day 1 declining to N = 1 on day 134) and C (eggs removed and replaced; light/dark gray bars, N = 36 on day 1 declining to N = 1 on day 148) queens of *Bombus terrestris* over the course of the experiment. Bar height: number of queens alive on each day in each treatment; pale relative to dark colour (within each bar): proportion of queens that showed a response to disturbance (became active/increased speed after the colony was moved) to queens that showed a non-response to disturbance (did not become active/increase speed after the colony was moved) on each day in each treatment. The proportion of responses to disturbance declined significantly with time for both treatments (binomial glmm: $b = -1.7963$, $SEb = 0.307$, $z = -5.856$, $p < 0.001$). However, there was no significant difference in queens' response to disturbance between R and C colonies (see Supplementary Results).

**Figure S9**

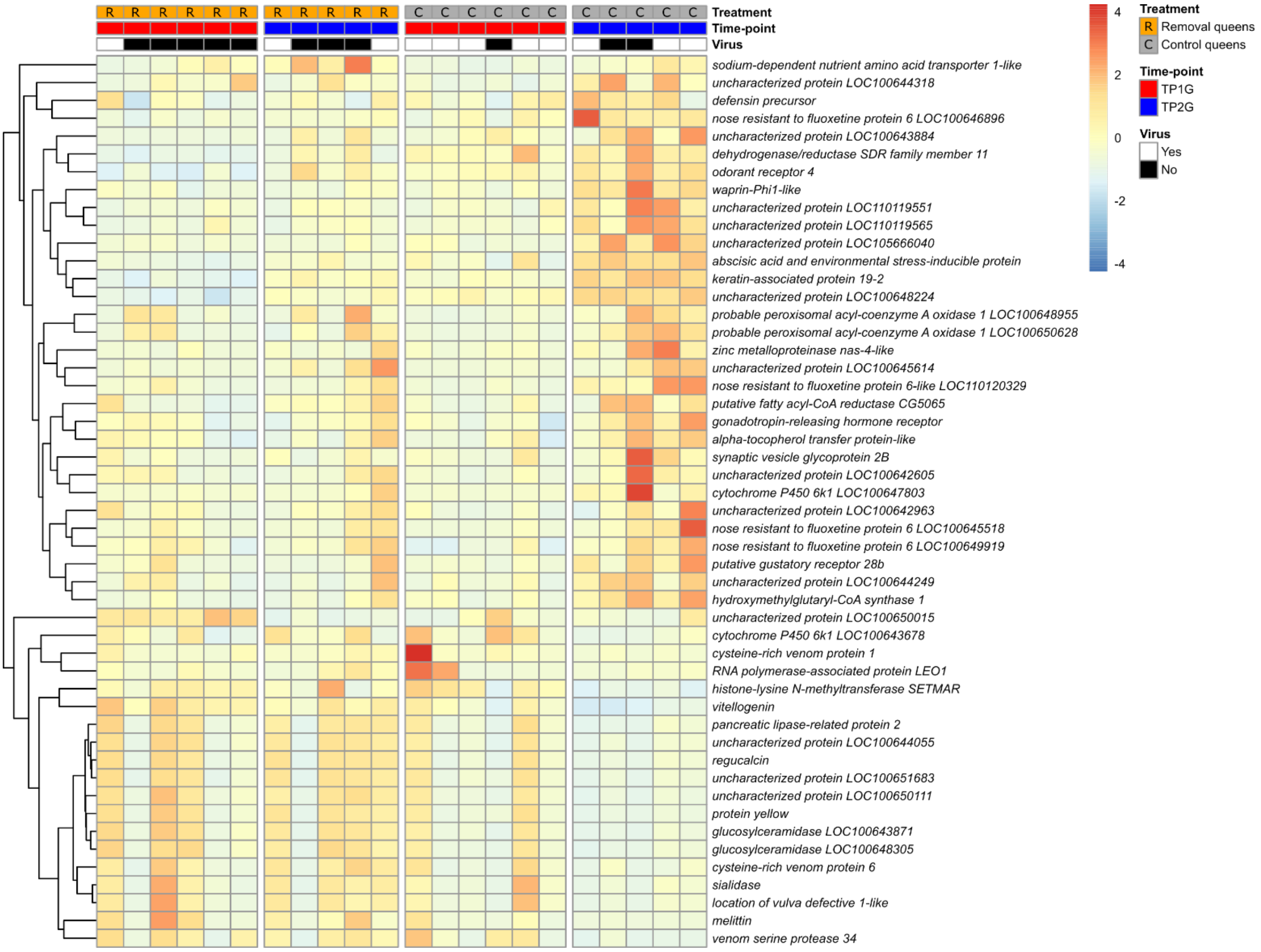

Figure S9. Gene expression differences from mRNA-seq libraries prepared from single brain samples from R (eggs removed) and C (eggs removed and replaced) *Bombus terrestris* queens. Differences expressed in a heatmap showing relative changes in gene expression ($log_2$ fold change) within each gene for the 50 most highly differentially expressed genes (DEGs) (out of 836 DEGs in total), with each row representing an individual gene and each column representing a biological replicate of the mRNA-seq data. Vertical breaks separate the two treatments (R (orange) and C (gray)) and the two time-points (TP1G (red) and TP2G (blue)). The presence of virus reads in the mRNA-seq library is annotated by white (Yes - present) and black (No – absent) bars. The dendrogram at left groups genes that cluster according to their gene expression patterns.

**Figure S10**

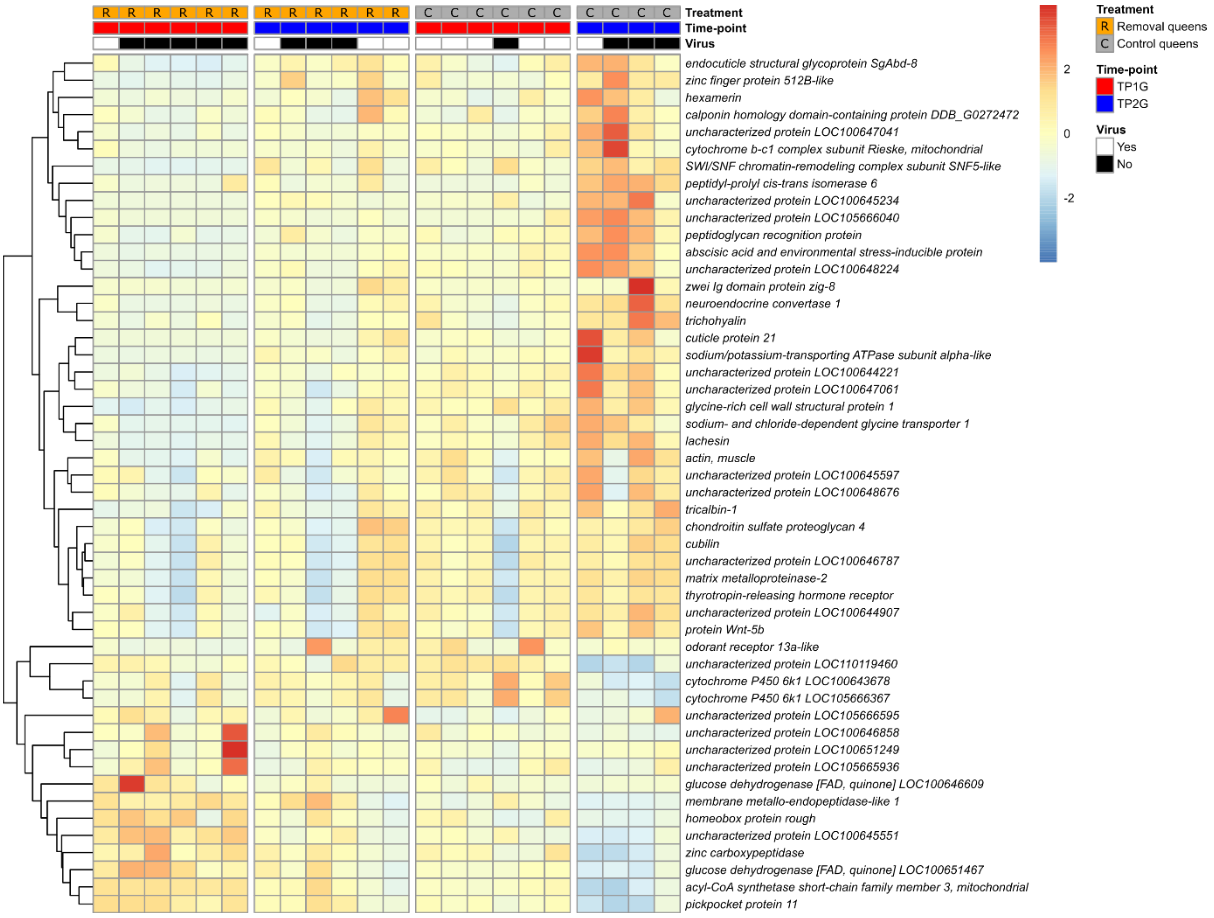

Figure S10. Gene expression differences from mRNA-seq libraries prepared from single fat body samples from R (eggs removed) and C (eggs removed and replaced) *Bombus terrestris* queens. Differences expressed in a heatmap showing relative changes in gene expression ($log_2$ fold change) within each gene for the 50 most highly differentially expressed genes (DEGs) (out of 2,572 DEGs in total), with each row representing an individual gene and each column representing a biological replicate of the mRNA-seq data. Vertical breaks separate the two treatments (R (orange) and C (gray)) and the two time-points (TP1G (red) and TP2G (blue)). The presence of virus reads in the mRNA-seq library is annotated by white (Yes - present) and black (No – absent) bars. The dendrogram at left groups genes that cluster according to their gene expression patterns.

**Figure S11**

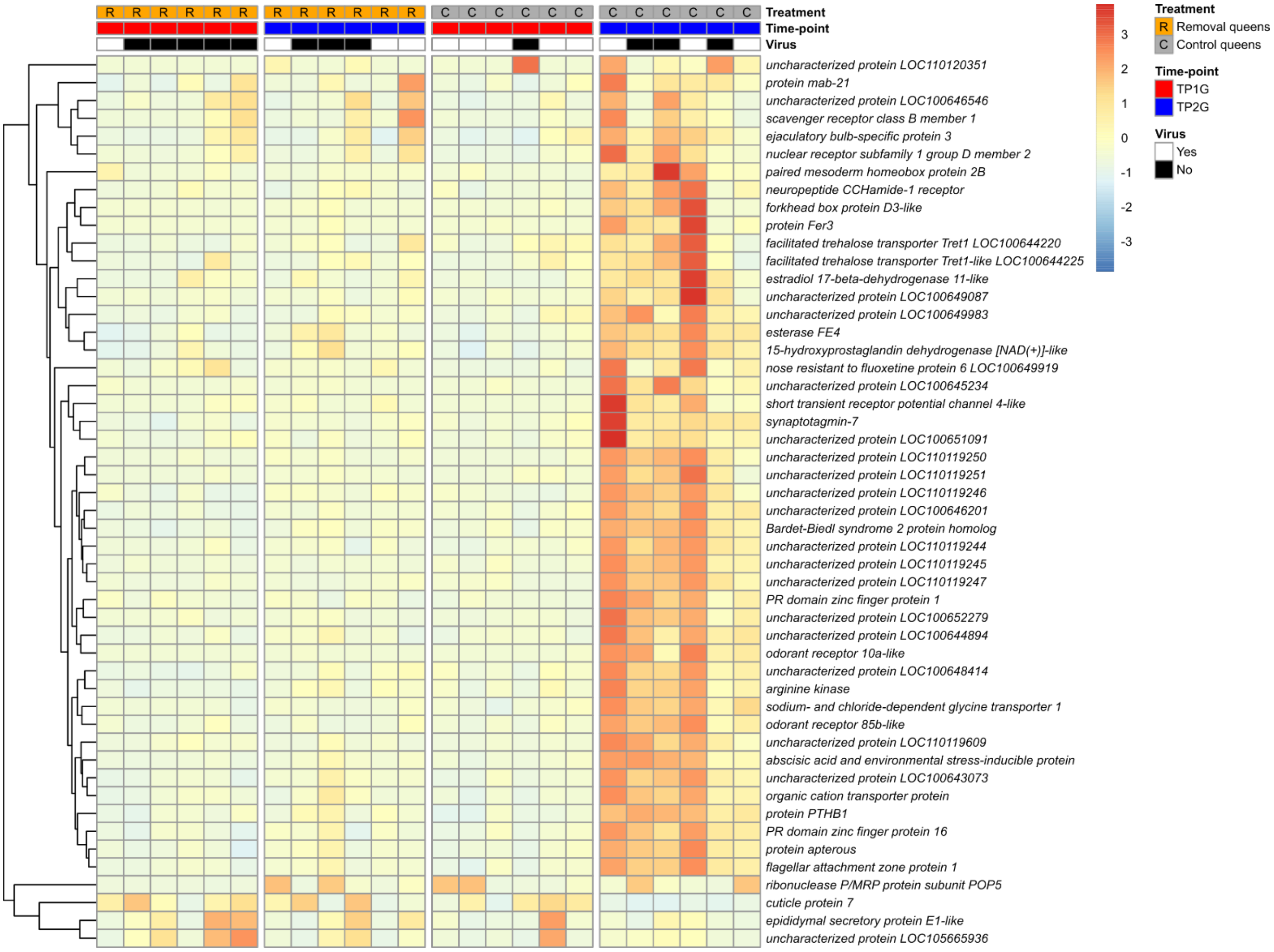

Figure S11. Gene expression differences from mRNA-seq libraries prepared from single ovaries samples from R (eggs removed) and C (eggs removed and replaced) *Bombus terrestris* queens. Differences expressed in a heatmap showing relative changes in gene expression ($log_2$ fold change) within each gene for the 50 most highly differentially expressed genes (DEGs) (out of 6,437 DEGs in total), with each row representing an individual gene and each column representing a biological replicate of the mRNA-seq data. Vertical breaks separate the two treatments (R (orange) and C (gray)) and the two time-points (TP1G (red) and TP2G (blue)). The presence of virus reads in the mRNA-seq library is annotated by white (Yes - present) and black (No – absent) bars. The dendrogram at left groups genes that cluster according to their gene expression patterns.

**Figure S12**

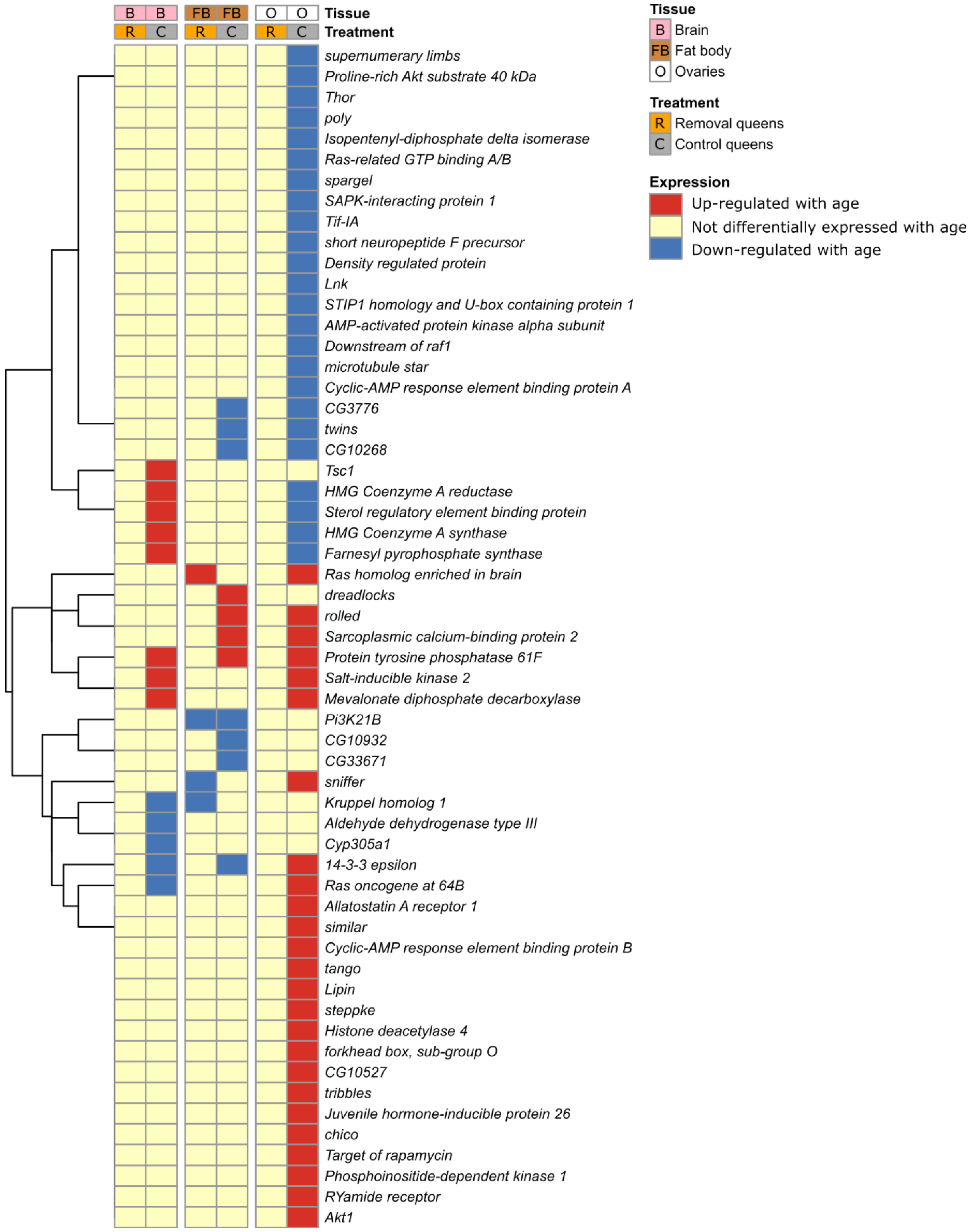

Figure S12. Results of comparison of age-related genes in *Bombus terrestris* queens and genes in the TI-J-LiFe network. Each row represents an individual gene from *Drosophila*

*melanogaster* described as a component of the TI-J-LiFe network by Korb et al. (2021) that has a single-copy orthologue in *B. terrestris* . Each column shows the age-related expression status of the focal genes in a given treatment and tissue in *B. terrestris* queens in the current study. Vertical breaks separate the three tissues (brain, fat body, and ovaries). The dendogram at left groups genes that cluster according to their gene expression patterns.

**Figure S13**

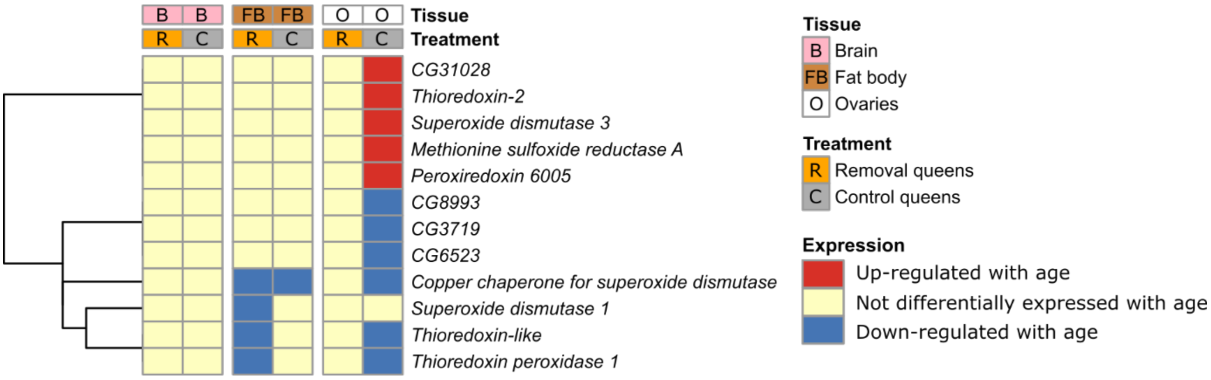

Figure S13. Results of comparison of age-related genes in *Bombus terrestris* queens and genes in the enzymatic antioxidant network. Each row represents an individual gene from *Drosophila melanogaster* known to be a component of the enzymatic antioxidant network described by Kramer et al. (2021) that has a single-copy orthologue in *B. terrestris*. Each column shows the age-related expression status of the focal genes in a given treatment and tissue in *B. terrestris* queens in the current study. Vertical breaks separate the three tissues (brain, fat body, and ovaries). The dendogram at left groups genes that cluster according to their gene expression patterns.

**Figure S14**

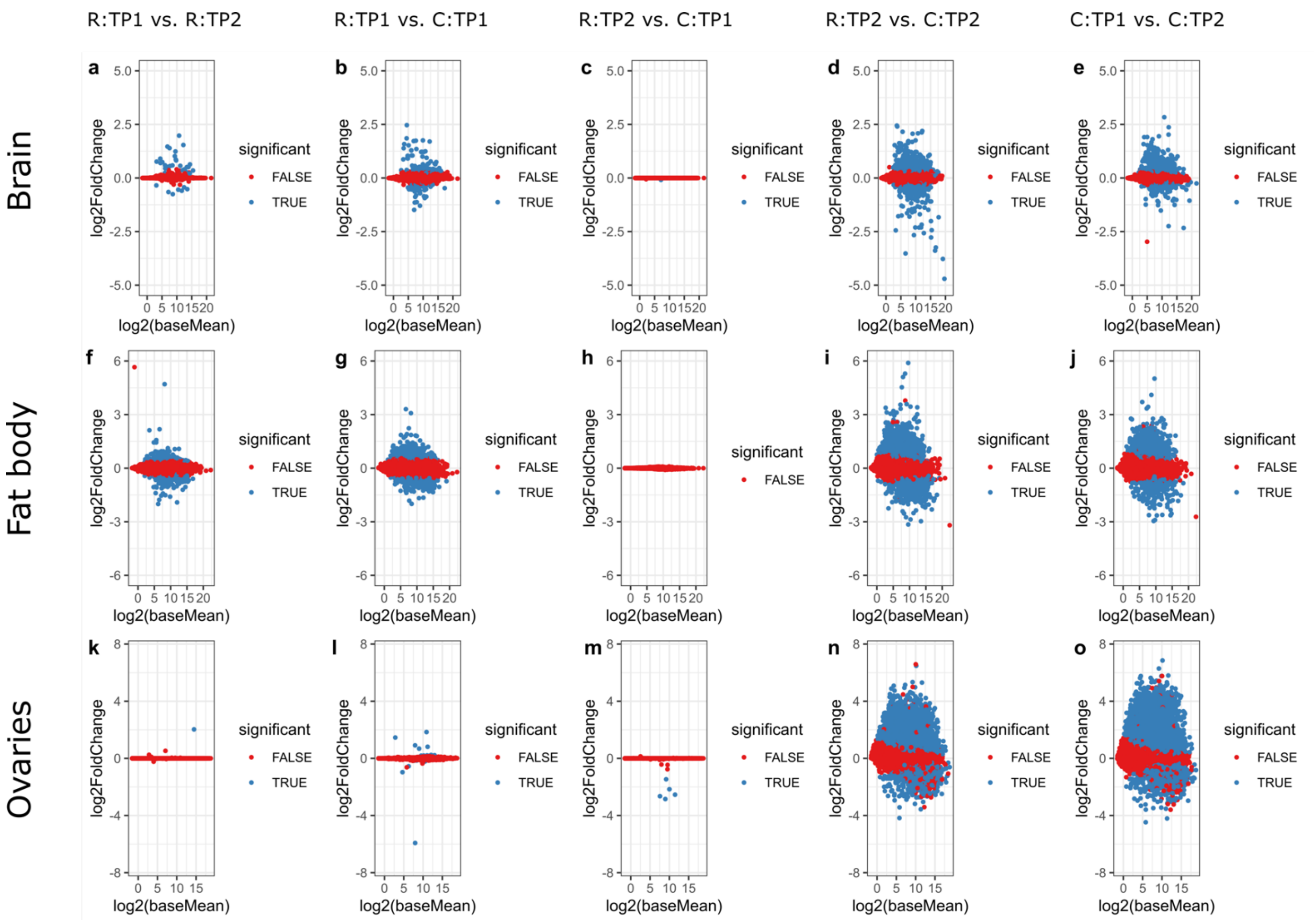

Figure S14. MA-plots comparing gene expression between pairs of treatments (R, eggs removed; C, eggs removed and replaced) and time-points (TP1 and TP2) from mRNA-seq libraries from three tissues of *Bombus terrestris* queens. X axis: $log_2$ value of the average expression of each gene in the experiment; Y axis: $log_2$ value of the fold-change of expression between two samples (i.e. $log_2$ of the ratio between expression of one sample and the other sample). **a-e,** brain; **f-j,** fat body; and **k-o,** ovaries. **a, f,** and **k**: comparisons between R:TP1 and R:TP2; **b, g** and **l**: comparisons between R:TP1 and C:TP1; **c, h** and **m**: comparisons between R:TP2 and C:TP1; **d, i** and **n**: comparisons between R:TP2 and C:TP2; **e, j** and **o**: comparisons between C:TP1 and C:TP2. Blue points ('true'): differentially expressed genes (DEGs); red points ('false'): genes that are not differentially expressed, based on DESeq2 analysis. N = 6 for all treatments/time-points except brain R:TP2 and C:TP2 (N = 5), and fat body C:TP2 (N = 4).

**Figure S15**

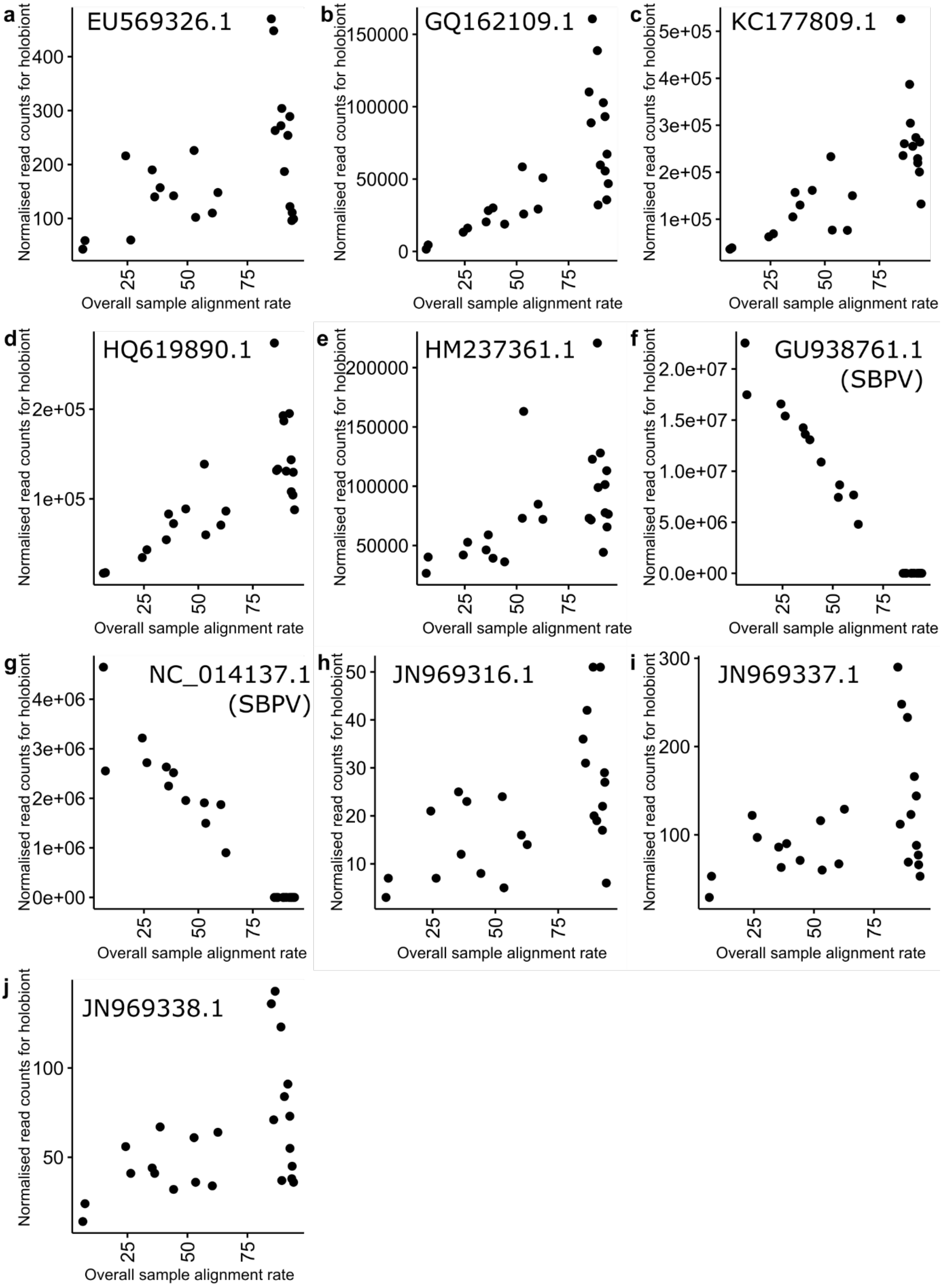

Figure S15. The relationship between presence of *Apis mellifera* holobiont sequences and alignment to the *Bombus terrestris* genome for fat body mRNA-seq libraries from the current study. The 10 sequences shown are those with >500 normalised counts in total from Kallisto across all the libraries from a tissue, in at least 2 out of the 3 tissues (brain, fat body, and ovaries). **a-j,** Scatterplots show fat body mRNA-seq libraries from the current study (N = 24) with normalised read counts from Kallisto for an *Apis mellifera* holobiont sequence (GenBank accession given at the top of the panel), plotted against the overall percentage alignment of the mRNA-seq library to the *Bombus terrestris* genome using HISAT2. Holobiont sequences in panels **f** and **g** are GenBank accessions for slow bee paralysis virus (SBPV).

**Figure S16**

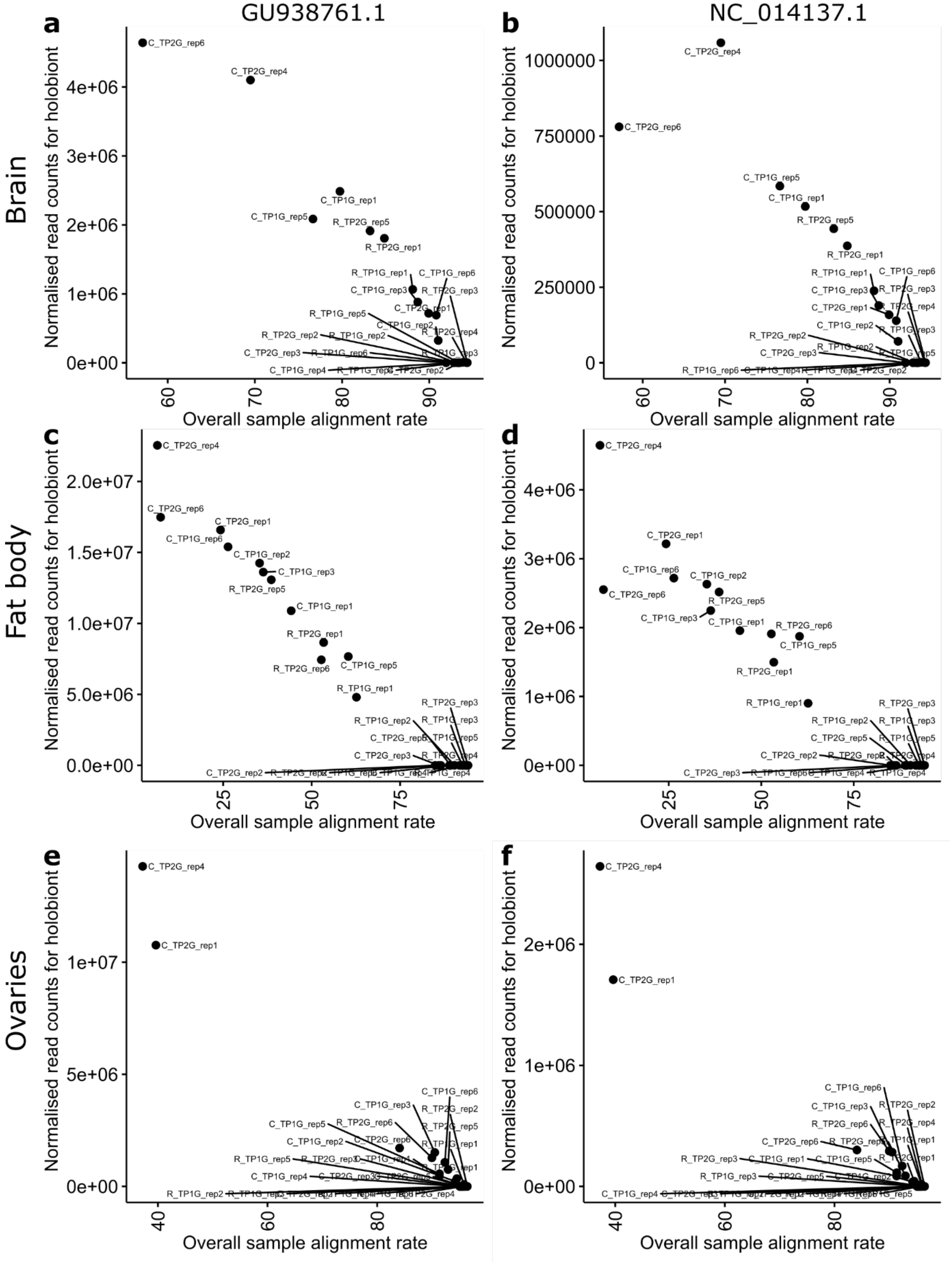

Figure S16. The relationship between presence of slow bee paralysis virus (SBPV) sequences and alignment to the *Bombus terrestris* genome for mRNA-seq libraries from the current study. Scatterplots show mRNA-seq libraries from the current study with normalised read counts from Kallisto for slow bee paralysis sequences (**a, c, e,** GU938761.1 or **b, d, f,** NC_014137.1) plotted against the overall percentage alignment of the mRNA-seq library to the *Bombus terrestris* genome using HISAT2 for libraries from **a, b,** brain, **c, d,** fat body or **e, f,** ovaries. Libraries are labelled with the library name (with, if required, a black line connecting the library name and the relevant point). Library names are in the format, treatment_time-point_biological replicate. R, removal queens (eggs removed); C, control queens (eggs removed and replaced); TP1G, time-point 1; TP2G, time-point 2; rep1, biological replicate 1. For the mRNA-seq libraries: Brain, N = 22; Fat body, N = 24; Ovaries, N = 24.

**Figure S17**

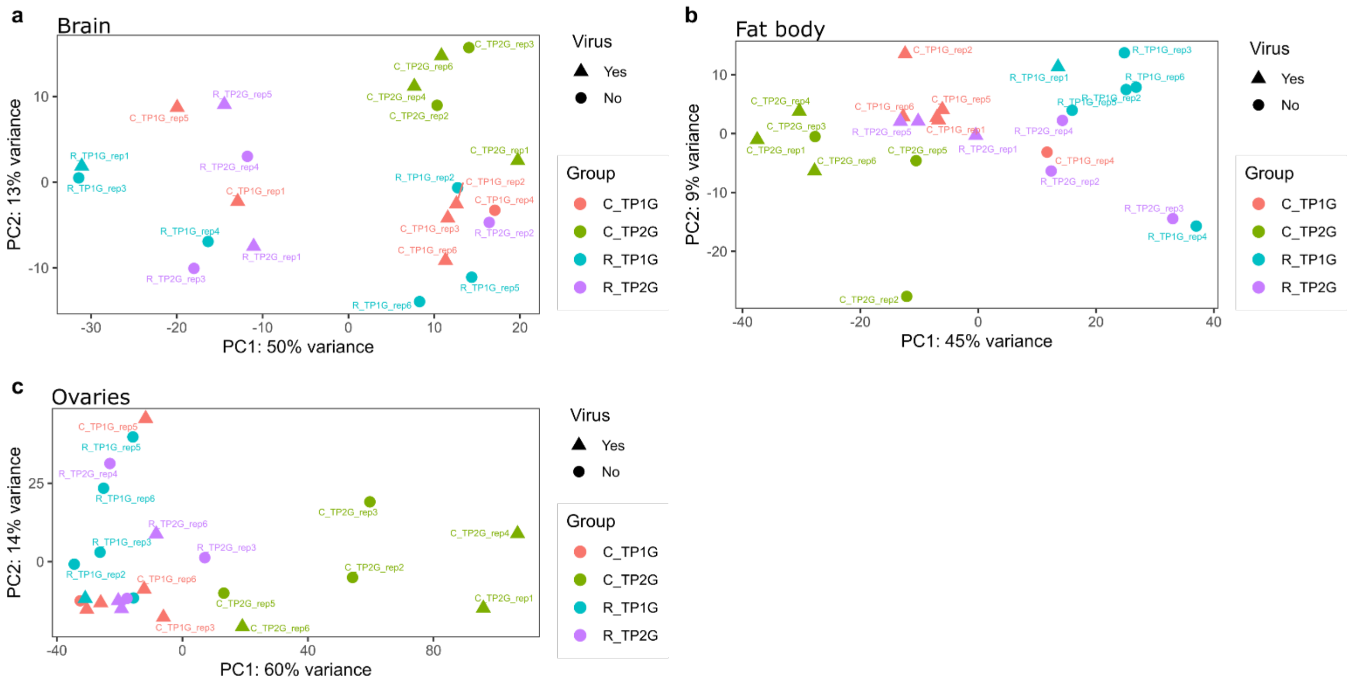

Figure S17. PCA plots showing that mRNA-seq libraries do not cluster by presence of slow bee paralysis virus (SBPV) reads. Principal component analysis (PCA) plots for the top 2000 most highly expressed genes isolated from mRNA-seq libraries from single *Bombus terrestris* queens from **a,** brain **b,** fat body and **c,** ovaries. **a, b, c,** Axes represent principal components. Individual points of the same colour indicate biological replicates of the same group. Libraries where large numbers of SBPV reads were present are denoted with triangles (yes), whereas libraries without large numbers of SBPV reads are denoted with circles (no). Libraries are labelled with the library name (with, if required, a line connecting the library name and the relevant point). Library names are in the format, treatment_time-point_biological replicate. Group names in the format treatment_time-point. R, removal queens (eggs removed); C, control queens (eggs removed and replaced); TP1G, time-point 1; TP2G, time-point 2; rep1, biological replicate 1. For the mRNA-seq libraries: **a,** Brain, N = 22; **b,** Fat body, N = 24; **c,** Ovaries, N = 24.

**Figure S18**

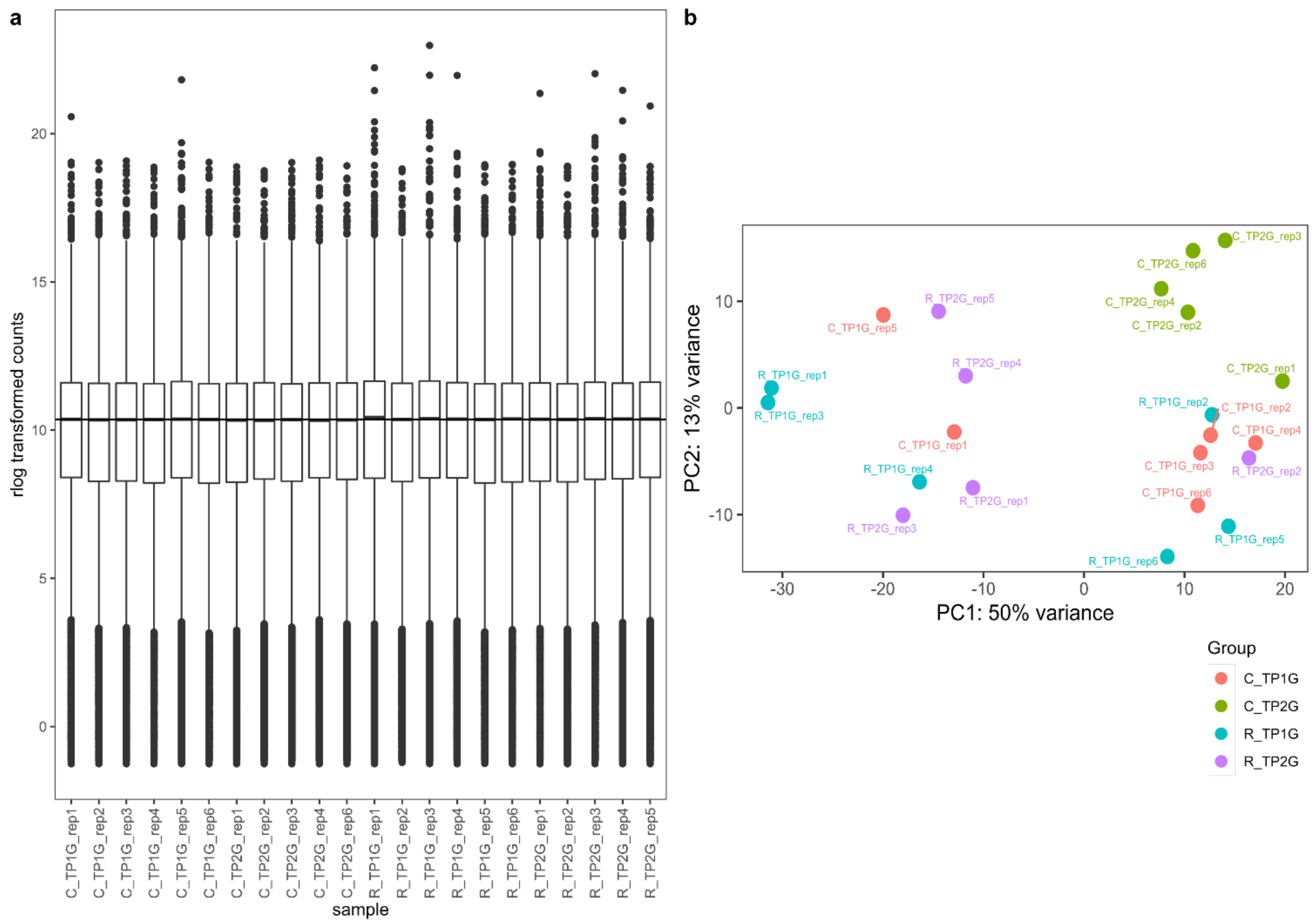

Figure S18. Exploratory plots from the differential expression analysis of mRNA-seq libraries from brain of single *Bombus terrestris* queens. **a,** Normalisation boxplots of the rlog-transformed value of mRNA-seq expression for genes in each library. Boxplots show median, interquartile range, 10th and 90th percentile. Horizontal line represents the median value of the transformed counts. **b,** Principal component analysis (PCA) plot of the top 2,000 most highly expressed genes isolated from mRNA-seq libraries in brain. Axes represent principal components. Individual points of the same colour indicate biological replicates of the same group. Libraries are labelled with the library name (with, if required, a line connecting the library name and the relevant point). **a, b,** Library names are in the format, treatment_time-point_biological replicate. Group names in the format treatment_time-point. R, removal queens (eggs removed); C, control queens (eggs removed and replaced); TP1G, time-point 1; TP2G, time-point 2; rep1, biological replicate 1. Brain mRNA-seq libraries: N = 22.

**Figure S19**

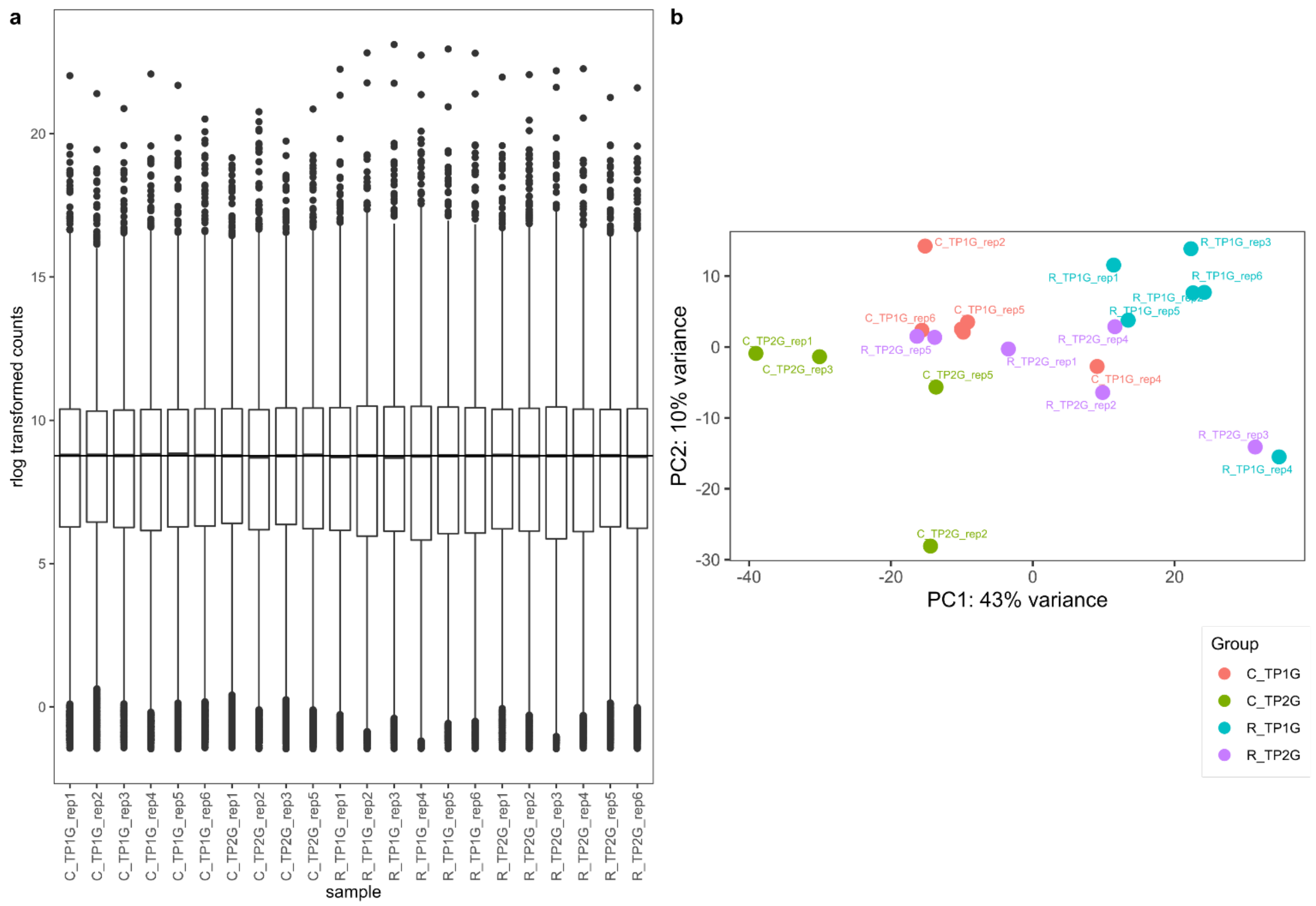

Figure S19. Exploratory plots from the differential expression analysis of mRNA-seq libraries from fat body of single *Bombus terrestris* queens. **a,** Normalisation boxplots of the rlog-transformed value of mRNA-seq expression for genes in each library. Boxplots show median, interquartile range, 10th and 90th percentile. Horizontal line represents the median value of the transformed counts. **b,** Principal component analysis (PCA) plot of the top 2000 most highly expressed genes isolated from mRNA-seq libraries in fat body. Axes represent principal components. Individual points of the same colour indicate biological replicates of the same group. Libraries are labelled with the library name. **a, b,** Library names are in the format, treatment_time-point_biological replicate. Group names in the format treatment_time-point. R, removal queens (eggs removed); C, control queens (eggs removed and replaced); TP1G, time-point 1; TP2G, time-point 2; rep1, biological replicate 1. Fat body mRNA-seq libraries: N = 22.

**Figure S20**

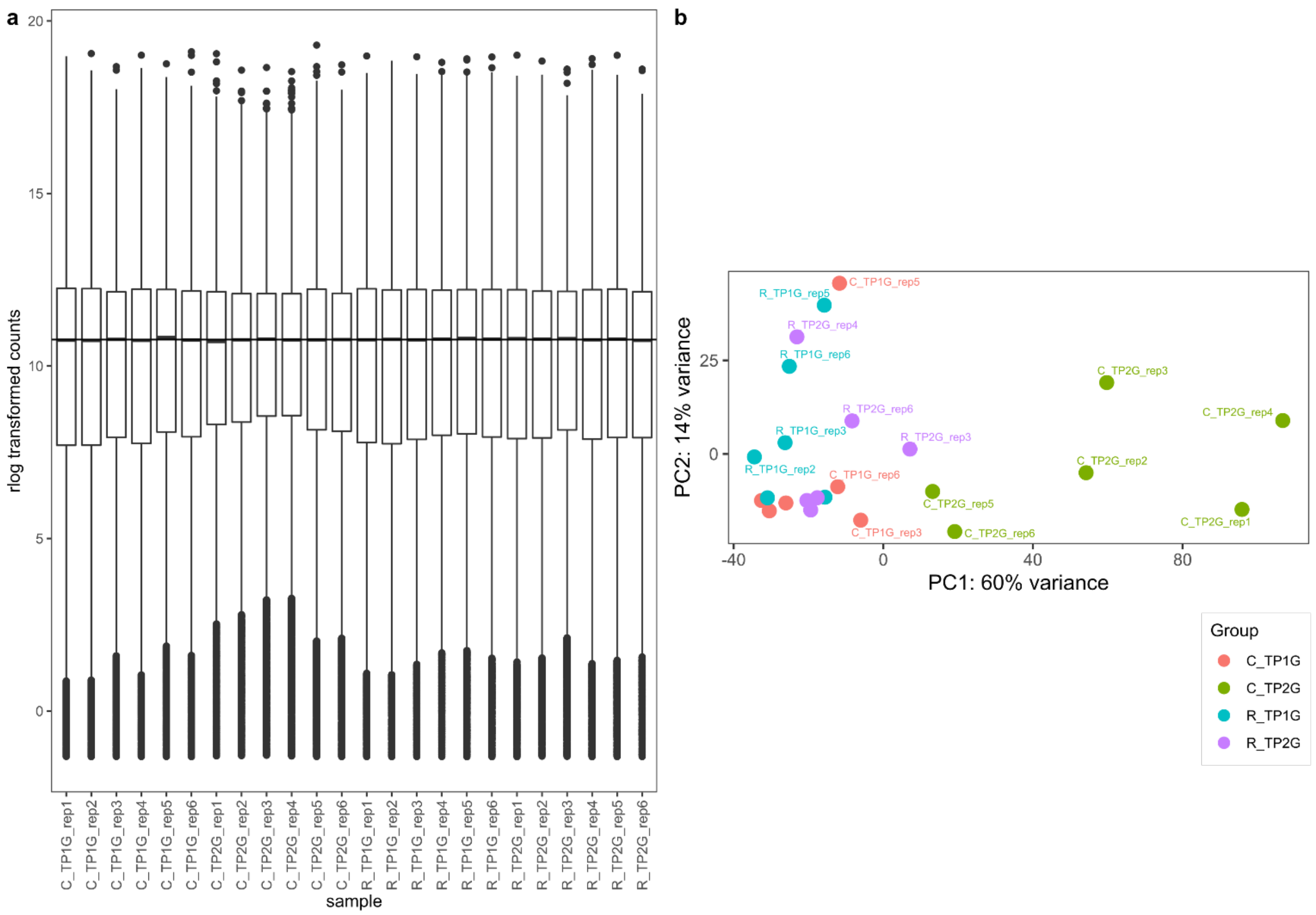

Figure S20. Exploratory plots from the differential expression analysis of mRNA-seq libraries from ovaries of single *Bombus terrestris* queens. **a,** Normalisation boxplots of the rlog-transformed value of mRNA-seq expression for genes in each library. Boxplots show median, interquartile range, 10th and 90th percentile. Horizontal line represents the median value of the transformed counts. **b,** Principal component analysis (PCA) plot of the top 2000 most highly expressed genes isolated from mRNA-seq libraries in ovaries. Axes represent principal components. Individual points of the same colour indicate biological replicates of the same group. Libraries are labelled with the library name. **a, b,** Library names are in the format, treatment_time-point_biological replicate. Group names in the format treatment_time-point. R, removal queens (eggs removed); C, control queens (eggs removed and replaced); TP1G, time-point 1; TP2G, time-point 2; rep1, biological replicate 1. Ovaries mRNA-seq libraries: N = 24.
